# HIF-1α+ CD4 T cells coordinate a tissue resident immune cell network in the lung

**DOI:** 10.1101/2025.06.13.659476

**Authors:** Jean de Lima, Nivedya Swarnalekha, Ewelina Bartoszek, Ludivine C. Litzler, Claire E. Depew, Mara Esposito, Maike Erber, Marco Künzli, Anukul T. Shenoy, Ananda W. Goldrath, Jie Sun, David Schreiner, Carolyn G. King

**Author notes:** Corresponding author: Carolyn King Department of Biomedicine University of Basel Mattenstrasse 28, CH-4058 Basel, Switzerland.

## Abstract

A deeper understanding of how tissue localized immune cells arise and function is critical for developing mucosal vaccines. Currently, there are no murine models that specifically target tissue T cells while leaving their lymphoid counterparts untouched. Here we leverage the observation that during influenza infection, HIF-1α regulatory activity is higher in the lung compared to lymph node CD4 T cells. Inducible deletion of *Hif1a* in CD4 T cells, at the onset of its activity in the lung, reduces the tissue resident T cell compartment with minimal impact on peripheral immunity. HIF-1α-active CD4 T cells occupy the border of tertiary lymphoid structures, where they coordinate an IL-21-dependent network of spatially co-localized immune cells including macrophages, NK cells and IgA+ B cells. A similar HIF-1α-dependent network is engaged in a lung adenocarcinoma model, highlighting a broader role for HIF-1α+ CD4 T cells in integrating protective immunity during infection and cancer.

## INTRODUCTION

T cell responses are initiated in draining lymphoid organs, where T cells recognize cognate antigen presented by antigen presenting cells. Following clonal expansion, T cells migrate to non-lymphoid tissues, where they differentiate into a heterogeneous array of effectors. This directional sequence of events, in which adaptive immunity originates in the lymph node and is executed in the tissue, is referred to as an “inside-out” immune response^1^. Ongoing inflammation in the tissue can trigger the formation of tertiary lymphoid structures (TLS), composed of T cells, B cells, dendritic cells, macrophages and stromal cells^2^. TLS are elicited in response to infection, autoimmunity and cancer and may play host protective or pathologic roles depending on the immunologic context. TLS have a well known role in supporting mucosal antibody production and have also been identified as sites for de novo T cell recruitment and priming, capable of independently organizing an “inside-out” immune response within the tissue^3^. However, the signals regulating T cell diversification within the TLS microenvironment remain poorly understood.

T cell diversification is governed by a variety of inputs including antigen receptor engagement, cytokines, hypoxia and nutrient availability, all of which are dynamically regulated in time and space. The distribution of these signals in and outside of the TLS likely plays a central role in directing tissue T cell fate. For example, influenza infection elicits the development of heterogeneous lung resident CD4 T cells including TLS-localized T resident helper (TRH) cells and Th1 cells which are distributed throughout the tissue^4–6^. TRH cells, similar to T follicular helper (Tfh) cells in lymphoid organs, require intrinsic Bcl6 expression, support local IgG antibody responses and retain the capacity to generate diverse secondary effectors^4,7,8^. TRH cells can be further divided into two distinct subsets regulated by the transcription factors Bcl6 and HIF-1α, respectively. While the role of Bcl6 in T cells has been extensively documented, the role of HIF-1α is more controversial^9^.

Genetic disruption of HIF-1α in T cells leads to pleiotropic effects on Th17, Treg and Th1 cell differentiation, with outcomes that are strongly influenced by microenvironmental cues^10–15^. In lymphoid organs, HIF-1α has been described to either restrain or promote TFH cell development^16–19^. In the tissue, HIF-1α stabilization promotes a residency gene signature and anti-tumor effector function in CD8 T cells but paradoxically impairs the accumulation of lung resident CD4 T cells, leading to enhanced disease susceptibility in a mouse model of tuberculosis infection^20–23^. One possible explanation for these discrepancies is that the genetic models used in prior studies ablated or stabilized HIF-1α prior to T cell priming. This prevents an assessment of whether HIF-1α has distinct effects at later time points or within specific tissue niches. Importantly, HIF-1α can be induced by exposure to antigen, TGFβ and hypoxia, all of which are implicated in the formation of both tissue resident T cells and TLS^21,24–28^. However, the spatial relationship between HIF-1α–activating signals and TLS, as well as their influence on the diversity and function of tissue-resident CD4 T cells, remain unclear.

To address this, we used spatial transcriptomics to examine the lymphocytes localized within and outside TLS during influenza infection. Strong T cell - B cell interaction and Tfh gene signatures were concentrated in the TLS core, while gene signatures for T cell receptor signaling and hypoxia extended into the TLS periphery, where a subset of HIF-1α-active CD4 T cells was also detected. Inducible deletion of HIF-1α, allowing temporal control of gene ablation after priming, revealed its central role in stabilizing the lung resident CD4 T cell compartment with negligible effect on peripheral T cell responses. Unexpectedly, HIF-1α+ CD4 T cells are responsible for coordinating a network of diverse immune responses including replenishment of alveolar macrophages, accumulation of lung resident NK cells and maintenance of influenza specific IgA antibody titers in the respiratory tract. These seemingly disparate responses converge upon a requirement for IL-21 provided by HIF-1α+ CD4 T cells.

## RESULTS

### Spatially structured immune cell interactions in lung TLS

To investigate the cellular architecture of TLS in an infection context we performed spatial transcriptomic profiling (Xenium, 10x Genomics) on murine lung tissue 14 days after influenza infection. TLS structures were clearly identifiable as high density regions of the B cell-specific Ig-beta gene *Cd79b* that were consistent with H&E staining (**Figure 1A**). Accordingly, annotated B cells populated the TLS core while peripheral regions were enriched for transcripts from dendritic cells, macrophages, T cells and plasma cells, forming a layer around the B cell-dominated core **(Figure 1B and C**). In order to link immune cell positioning to known lymphoid organization cues, we assessed chemokine and receptor expression as a function of distance from the edge of TLS (**Figure 1D**). The TLS core areas were enriched for transcripts associated with B cell follicles in SLO including *Cxcl13* and its receptor *Cxcr5*, as well as *Cxcr4* and *Ccr6* which mark B cell memory precursors^29,30^. Increased expression of *Ccr7, Ccl19, Cxcr3, Cxcl9 and Cxcl10* at the boundary of the B cell core suggests this region may be a site for the recruitment and retention of CD4 T cells and resident memory B cells^31–33^. Chemokine transcripts found primarily outside of the B cell core included *Ccr2*, *Ccl21a/b*, and *Cxcr6* with its ligand *Cxcl16*^34,35^. Spatially distinct expression of PD-1 ligands was also observed: *Cd274* (encoding PD-L1) was more prominent in macrophages and B cells outside TLS, while B cells inside TLS expressed more *Pdcd1lg2* (encoding PD-L2) (**Figure 1E and S1A**). This parallels findings in lymph nodes where PD-L1 from bystander B cells restricts Tfh cell access to follicles unless overridden by ICOS signaling^36^. Interestingly, peak expression of receptors and their ligands was not always colocalized, suggesting dynamic migration processes or the predominance of other signals. Examples include *Cxcr4* - *Cxcl12* and *Ccr7* -*Ccl21a/b*.

**Figure 1.**
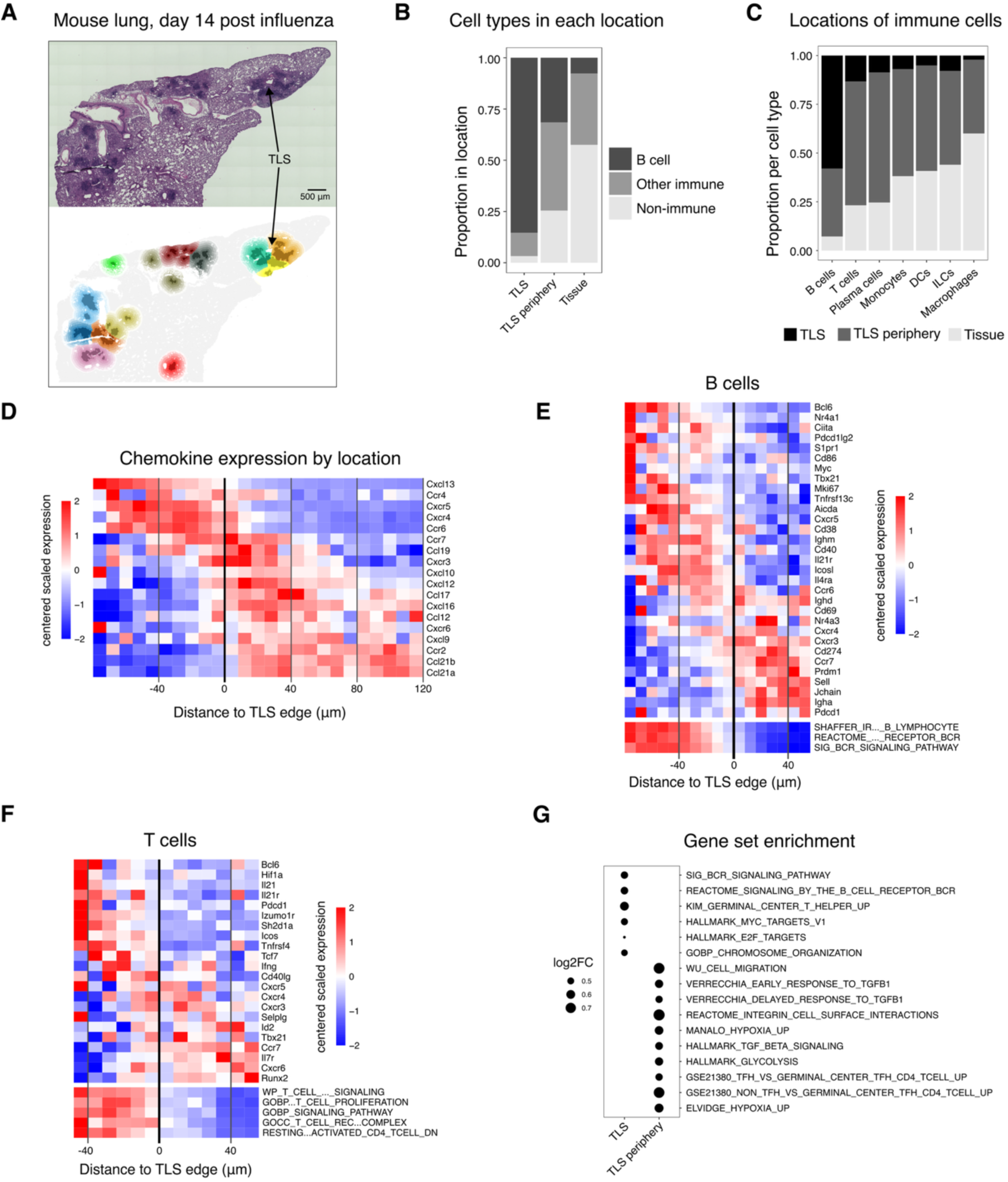
Spatially structured immune cell interactions in lung TLS. (A-G) Xenium spatial transcriptomic data, mouse lung day 14 post influenza, one section (*n* = 1 mouse). (A) H&E (top) and Xenium spatial transcriptomic transcriptome analysis (bottom) of mouse lung at 14 days post influenza. Bottom colored by *Cd79b* high TLS structure (dark) with surrounding periphery (faded). (B) Proportions of annotated cell types within each tissue location. (C) Proportions of tissue location within selected annotated cell types. (D-F) Centered scaled expression of genes or calculated MSigDB gene signature scores averaged in 8 μm bins by distance from the iBALT borders (*n* = 1 mouse). Rows are sorted using the maximum of a rolling average across 5 bins with a 1 bin step. The line at zero marks the iBALT edge; negative values are inside iBALT. Maximum and minimum values are limited to -2 or 2 respectively. (D) Chemokine-related genes on all cells. (E) Selected genes and gene set scores on B cells. (F) Selected genes and gene set scores on T cells. (G) Gene sets enrichment on all cells contrasting TLS core with TLS periphery. Statistical analyses: gene set enrichment analysis using limma camera, FDR < 0.05.

Building on these observations, we examined B and T cell gene expression patterns in the core and peripheral regions of the TLS. Core B cells resembled germinal center (GC) B cells expressing genes such as *Cd40, Il21r, Bcl6, and Cxcr5* (**Figure 1E**). B cells deepest inside the largest TLS expressed light zone genes (*Ciita, Pdcd1lg2, S1pr1*, *Cd86, and Myc)*, while those near the edge expressed dark zone associated genes (*Aicda, Mki67, Cxcr4,* and *Il4ra)* suggesting an anatomically inverted organization compared to GCs in SLO (**Figure 1E**)^30,31^. Outside the TLS core, B cells were enriched for memory-associated (*Cxcr3, Ccr6, Pdcd1, Nr4a3*) and plasma cell-associated (*Prdm1, Cxcr4, Jchain, Igha*) genes, with immunofluorescence staining confirming the location of IgA+ cells^37,38^ (**Figure 1E, Figure S1B and C**). T cells in the TLS core exhibited a Tfh phenotype (*Bcl6*, *Sh2d1a*, *Icos*, *Tcf7*, *Pdcd1*, and *Il21)* while T cells outside the core showed higher expression of *Cxcr3*, *Cxcr6*, *Tbx21*, *Selplg*, *Id2* and *Il7r*, consistent with type 1 immunity^4,39,40^ (**Figure 1F**). Gene set enrichment analysis corroborated these findings with elevated signatures for GC B cells (BCR signaling, cell cycle and *Myc*) and GC Tfh cells at the TLS core. Similar to what has been described at the T cell-B cell border in SLO, the outer TLS was also enriched for a non-GC Tfh signature, suggesting a functional division between Tfh cells located in the core compared to those in the periphery. Notably, signatures for hypoxia and TGFβ signaling, both known to support HIF-1α in T cells, were also present at the TLS periphery (**Figure 1F and G**)^18,19,21,41–43^. This pattern aligns with our previous finding that two Tfh like subsets - one enriched for Bcl6 activity and other for HIF-1α activity — emerge in the lung during influenza^4^. Taken together, TLS structures formed during influenza infection are spatially compartmentalized, with a central core enriched for GC Tfh cells and a peripheral zone containing non-GC Tfh cells. Increased hypoxia and TGFβ associated signaling at the TLS periphery suggest it may represent a distinct microenvironment that drives HIF-1α activity to uniquely shape T cell differentiation.

### Lung CD4 T cells increase HIF-1α activity during influenza infection

To understand the dynamics of *Hif1a* expression during influenza infection we re-analyzed single cell RNA sequencing (sc-RNAseq) data of influenza (NP) specific CD4 T cells isolated from the lung and draining lymph nodes at days 9, 14 and 30^4^. Transcriptional regulatory activity of HIF-1α was higher in lung CD4 T cells compared to those in the lymph node, increasing over time and peaking at day 30 (**Figure 2A**). To study this in more detail, we infected *Egln3*-YFP mice, which report the transcriptional activation of HIF-1α^41^. Consistent with the sequencing data, *Egln3*-YFP was progressively upregulated in both lung and lymph node CD4 T cells (**Figures 2B and S2A**). Sorted YFP+ TRH cells (folate receptor 4 (FR4)+, Egln3+) had enhanced gene expression of *Hif1a* compared to both YFP- TRH cells (FR4+, Egln3-) and tissue resident Th1 (TRM1, FR4- Egln3-) cells, validating the use of YFP as a readout for HIF-1α activity (**Figure S2B**). In the lung, *Egln3*-YFP expression was restricted to FR4 positive CD4 T cells, marking the TRH compartment (**Figure 2B**). Although the frequency of lung *Egln3-*YFP+ CD4 T cells increased until day 30, it sharply decreased at later time points, indicating that HIF-1α activity delineates a transient lung effector subset with Tfh cell characteristics (**Figure 2C**). In the draining lymph node, *Egln3*-YFP was primarily expressed by CXCR5+ Tfh cells **(Figure S2A**) and, in contrast to its transient expression in the lung, was maintained until at least 100 days after infection (**Figure 2C**). No appreciable *Egln3*-YFP was detected in CD4 T cells from the blood or in lung “circulating” T cells that stained positively for intravenous (iv) anti-CD45, suggesting that lung CD4 T cells independently upregulate HIF-1α activity in response to local cues (**Figure 2C**). Next, we used histology to assess where *Egln3*-YFP+ CD4 T cells are localized in the infected lung. Unlike Bcl6-high CD4 T cells which are centrally contained within iBALT^4^, *Egln3*-YFP+ CD4 T cells are more abundant at the iBALT perimeter (**Figure 2D**). The distinct positioning of these TRH subsets is in agreement with earlier observations that TRH cells with high Bcl6-regulatory activity have a transcriptional signature of lymphoid residency, while TRH cells with high HIF-1α regulatory activity are enriched for tissue residency genes^4^. To determine whether Egln3+ CD4 T cells in the lung depend on TLS, we treated influenza infected mice with LTβR-Fc to prevent TLS formation^42^. LTβR-Fc treatment led to a clear reduction of Egln3+ CD4 T cells in the lung while total numbers of influenza specific T cells were less affected (**Figures 2E and F, S2C and S2D**). Together these data reveal a transient subset of HIF-1α-active TRH cells that likely originate from local TLS.

**Figure 2.**
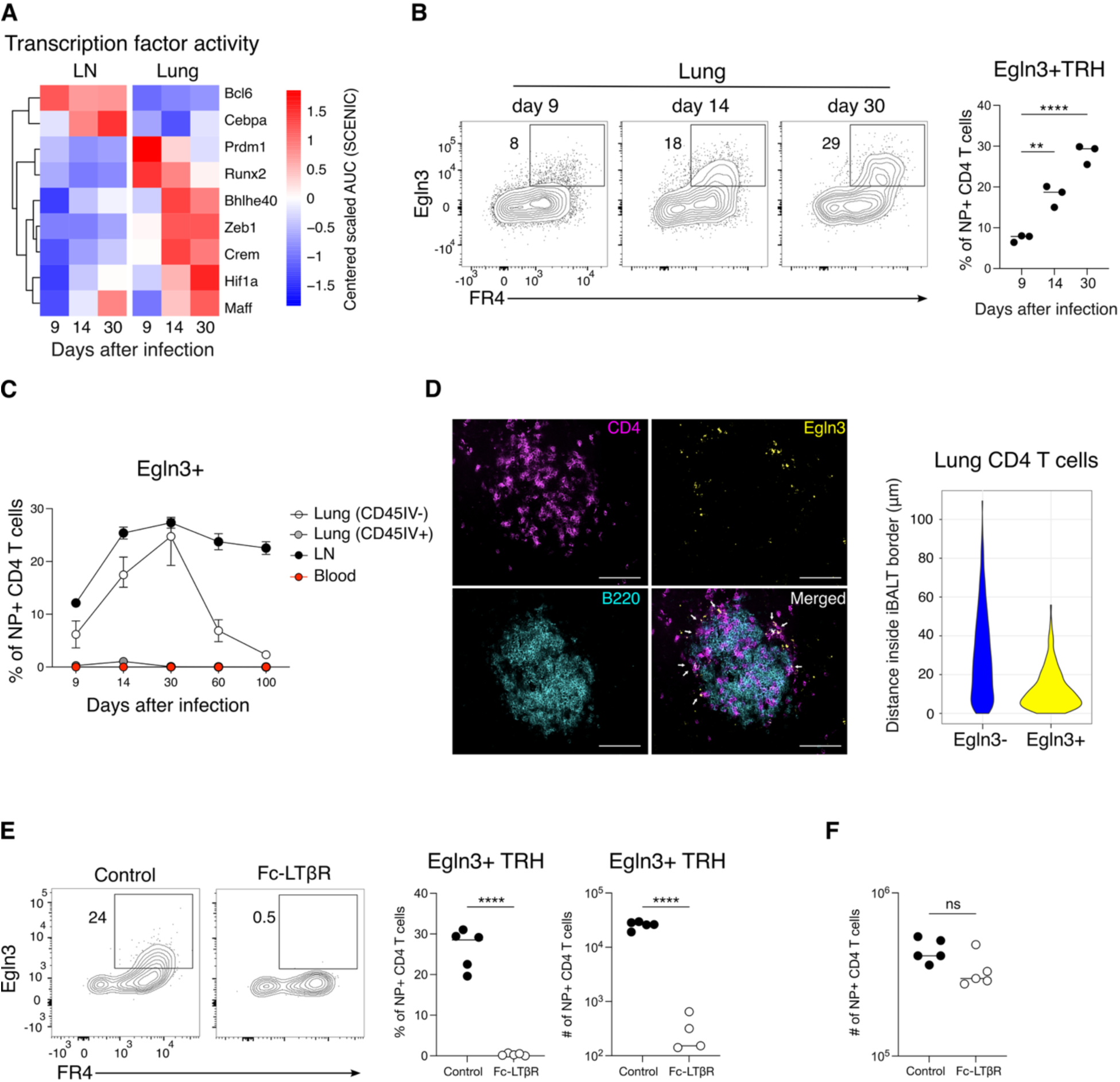
Lung CD4 T cells increase HIF-1α activity during influenza infection. (A) Centered, scaled SCENIC AUC of scRNA-seq data from NP+ CD4 T cells in the lung and lymph node (LN) (*n =* 3 mice). (B) Flow cytometry analysis showing frequency of lung NP-specific Egln3+ FR4+ (TRH) CD4 T cells from Egln3-YFP mice at 9, 14, and 30 days post-PR8 infection (mean ± SD, *n =* 3 mice). (C) Dynamics of Egln3-YFP+ NP+ CD4 T cells across lung resident, circulating, LN, and blood compartments (mean ± SD, *n =* 4 mice). (D) Confocal microscopy of iBALT showing Egln3+ and Egln3-CD4 T cells and distribution of distances (μm) of Egln3- and Egln3+ CD4 T cells from iBALT borders in 50 iBALT across mice using confocal images. Bars, 100µm (mean ± SD, *n =* 3 mice). (E) Frequency and numbers of NP-specific Egln3+ TRH cells in Egln3-YFP mice at day 20 post-PR8 infection, treated i.p. with isotype control or Fc-LTβR on days 9 and 14 (mean ± SD, *n =* 5 mice). (F) Numbers of resident NP-specific CD4 T cells in Egln3-YFP mice at day 20 post-PR8 infection, treated i.p. with isotype control or Fc-LTβR on days 9 and 14 (mean ± SD, *n =* 5 mice). Statistical significance determined by ANOVA (B) and t-tests (E). Adjusted p-values: *p<0.05, **p<0.005, ***p<0.0005, ****p<0.0001, ns: not significant. Data representative of two independent experiments.

### HIF-1α+ CD4 T cells produce IL-21 and support T cell residency in the lung

To specifically address the role of HIF-1α in lung CD4 T cells, we generated Hif1a^flox/flox^CD4^Cre-ERT2^ (Hif1a^iKO^) mice, enabling tamoxifen-inducible *Hif1a* deletion in CD4 T cells. Heterozygous control (Hif1a^flox/+^CD4^Cre-ERT2^, Hif1a^ctrl^) and Hif1a^iKO^ mice were infected with influenza and treated with tamoxifen starting at day 15 after infection (**Figure 3A**), allowing us to preserve HIF-1α activity during initial priming of CD4 T cells in lymph nodes. Five days after tamoxifen treatment, the total number of influenza specific lung resident CD4 T cells were decreased by approximately 10-fold in Hif1a^iKO^ mice, while neither circulating CD4 T cells in the lung nor Tfh cells in the draining lymph node were affected (**Figures 3B, S3A and S3B**). Unexpectedly, TRM1 cells, defined by the expression of P-selectin glycoprotein ligand-1 (PSGL1)^4^, were significantly decreased in Hif1a^iKO^ mice while the absolute number of TRH cells was comparable across the strains **(Figure S2C).** Consistent with limited effects on TRH cells, *Hif1a* deletion had no impact on influenza-specific IgG titers in the bronchoalveolar lavage fluid (BALF) or serum **(Figure S3D)**. Our earlier work demonstrated that similar to TFH cells in lymphoid organs, adoptively transferred TRH cells maintain differentiation plasticity while TRM1 cells are more terminally differentiated^4,8^. We therefore considered the possibility that HIF-1α, most highly expressed by TRH cells, is required to mediate their transition to TRM1 cells. In support of this idea, visualization of surface markers FR4 and CXCR6, expressed on TRH and TRM1 cells, respectively, revealed an intermediate subset of FR4+CXCR6+ T cells which was ablated following *Hif1a* deletion (**Figure 3C**). HIF-1α activity was also detected within the FR4+CXCR6+ compartment, whose numbers peaked at day 14 after infection (**Figures S3E and S3F**).

**Figure 3.**
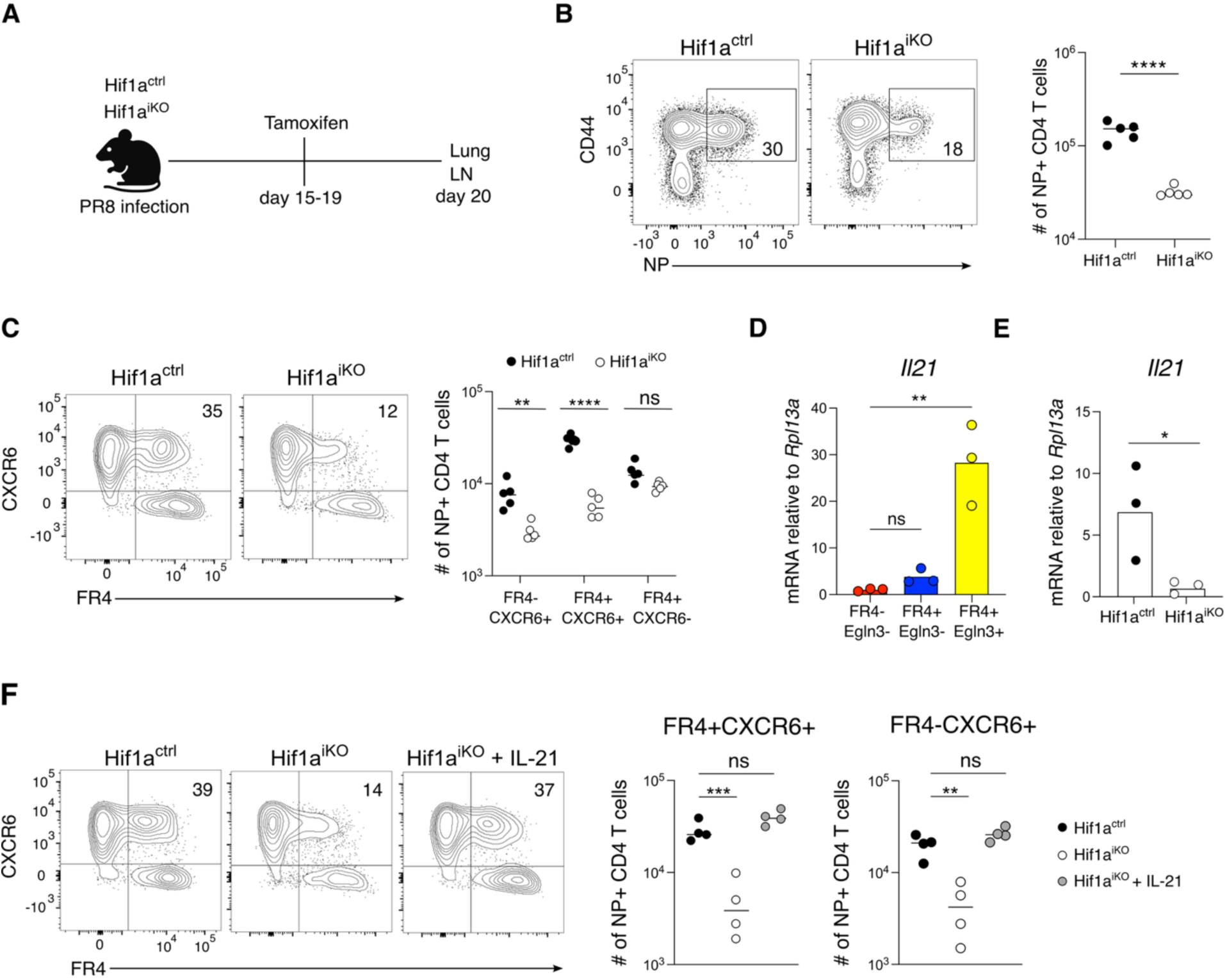
HIF-1α+ CD4 T cells produce IL-21 and support T cell residency in the lung. (A) Experimental design for (B-F). (B) Flow cytometry analysis and quantification of resident (ivCD45-) NP-specific CD4 T cells (mean ± SD, *n =* 5 mice). (C) Analysis and quantification of resident NP-specific CD4 T cell subsets in Hif1a^ctrl^ and Hif1a^iKO^ mice (mean ± SD, *n =* 5 mice). (D) *Il21* expression measured by TaqMan qPCR in sorted FR4-, FR4+ Egln3-, and FR4+ Egln3+ NP-specific CD4 T cells (mean ± SD, each dot represents 2 pooled mice). (E) Sorted FR4+ NP-specific CD4 T cells from Hif1a^ctrl^ and Hif1a^iKO^ mice (mean ± SD, each dot represents 2 pooled mice). (F) Quantification of FR4+CXCR6+ and FR4-CXCR6+ NP-specific CD4 T cells in Hif1a^ctrl^ and Hif1a^iKO^ mice following intratracheal IL-21 treatment (days 14–20) (mean ± SD, *n =* 4 mice). Statistical significance determined by t-tests (A), one-way ANOVA (D, F) or two-way ANOVA (C); adjusted p-values: *p<0.05, **p<0.005, ***p<0.0005, ****p<0.0001, ns: not significant. Data representative of two independent experiments.

We next addressed which HIF-1α derived signals might lead to upregulation of CXCR6 on lung TRH cells. At day 30 after influenza infection, approximately 50% of TRH cells actively transcribe IL-21, a cytokine previously reported to be decreased in HIF-1α deficient CD4 T cells cultured under Th17 polarizing conditions^4,13^. Similar to HIF-1α activity, IL-21 expression was detected in both FR4+CXCR6- and FR4+CXCR6+ T cells (**Figure S2G**). To investigate a linkage between HIF-1α activity and IL-21 production we assessed *Il21* transcripts in sorted *Egln3*-YFP+ TRH cells and in TRH cells isolated from Hif1a^iKO^ mice. *Il21* was more highly expressed by YFP+ TRH cells compared to YFP- TRH cells and TRM1 cells, and was strongly decreased in TRH cells following *Hif1a* deletion (**Figures 3D and 3E**). *Hif1α* deletion also led to decreased expression of PD-1 on lung NP+ CD4 T cells, in agreement with one study showing that exposure of CD4 T cells to IL-21 increases PD1 expression and another report showing that IL-21 deficient TFH cells express lower levels of *Pdcd1* transcript (**Figure S3H**)^43,44^. To further understand if local production of IL-21 contributes to the differentiation of lung CXCR6+ T cells, Hif1a^iKO^ mice were treated with intratracheal IL-21 concurrent with tamoxifen gavage. Provision of exogenous IL-21 was sufficient to restore the expression of both CXCR6 and PD1 on FR4+ and FR4- T cells in Hif1a^iKO^ mice (**Figure 3F and S3I**). Together these data raise the possibility that HIF-1α dependent IL-21 secretion by lung TRH cells supports CXCR6 upregulation and ng of tissue resident T cells outside of iBALT structures.

### HIF-1α+ CD4 T cells orchestrate lung immunity during influenza

The sole ligand for CXCR6 is the chemokine CXCL16, constitutively expressed by lung epithelial cells and macrophages^45,46^. Our secondary analyses of lung macrophage scRNAseq data revealed increased expression of both *Cxcl16* and *Il21r* two weeks after influenza infection, coinciding with the onset of HIF-1α activity and peak accumulation of FR4+CXCR6+ CD4 T cells in the lung (**Figure S4A, 2B and S3F**)^47^. In the spatial data, *Cxcl16* transcripts were abundantly expressed in the TLS periphery, where both *Cxcl16*+ and *Il21r*+ macrophages outnumbered their non-expressing counterparts (**Figures S4B and S4C**). Although *Il21r* transcripts primarily coincided with *Cd79b* expressed by B cells in the TLS core, they were also detectable in this peripheral area and could be assigned to a variety of cell types including T cells, monocytes and macrophages (**Figure S1A**). While *Izumor1r+Cxcr6+* T cells were more scarcely detected, they were concentrated in outer TLS, highlighting a potential role for these cells in supporting lung macrophage recovery after infection (**Figure S4D**).

Given the putative co-localization of HIF-1α-active CD4 T cells and CXCL16+ myeloid cells, we assessed the impact of *Hif1a* deletion in CD4 T cells on lung macrophages which are known to be transiently depleted and replenished during influenza infection^47–49^. In Hif1a^iKO^ mice, alveolar macrophages (AMs) were diminished and had decreased expression of MHC-II, CD86 and LFA-1 compared to Hif1a^ctrl^ mice, deficiencies which could be restored by exogenous IL-21 (**Figures 4A-C and S4E**). To understand if IL-21 is intrinsically required by lung macrophages, we generated irradiation chimeras transplanted with a mixture of wildtype and IL-21R^-/-^ bone marrow (**Figure S4F**). At baseline, reconstitution of the BALF was biased toward IL-21R^-/-^ immune cells, including both AMs and lymphocytes, while the lung was slightly better reconstituted by wildtype lymphocytes (**Figure S4F**). However, 14 days after influenza infection, IL-21R^-/-^ AMs in the BALF and monocyte derived macrophages in the lung were at a clear competitive disadvantage compared to wildtype cells (**Figure 4D, S4G and S4H**). Together these data indicate that macrophage intrinsic IL-21 sensing is required for full recovery of the lung macrophage compartment following infection.

**Figure 4.**
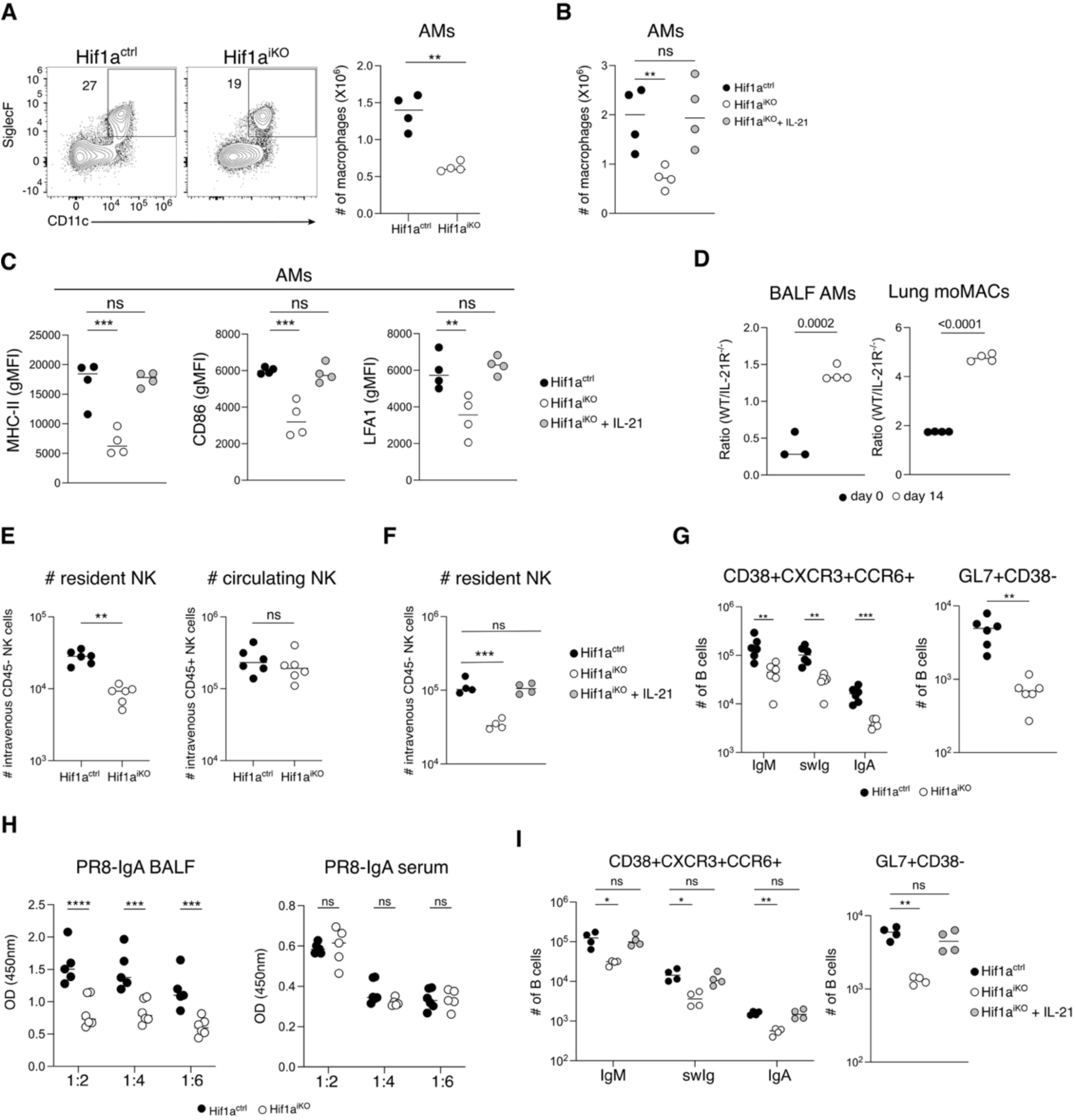
HIF-1α+ CD4 T cells orchestrate the lung immune response during influenza. (A) Representative flow cytometry plot and number of AMs from the lungs of Hif1a^ctrl^ and Hif1a^iKO^ mice (mean ± SD, *n =* 4 mice). (B) Numbers of AMs of Hif1a^ctrl^ and Hif1a^iKO^ mice following intratracheal IL-21 treatment (days 14–30) (mean ± SD, *n =* 4 mice). (C) MHCII, CD86, and LFA1 expression (gMFI) of lung AMs of Hif1a^ctrl^ and Hif1a^iKO^ mice following intratracheal IL-21 treatment (days 14–30) (mean ± SD, *n =* 4 mice). (D) Ratio of AMs in BALF and Lung monocyte-derived macrophages (moMACs) of chimeric mice (WT/IL-21R-/-) (mean ± SD, *n = 3-*4 mice). (E) Number of lung circulating and resident NK cells in Hif1a^ctrl^ and Hif1a^iKO^ mice following PR8 infection (mean ± SD, *n =* 6 mice). (F) Lung resident NK cell numbers in Hif1a^ctrl^ and Hif1a^iKO^ mice following intratracheal IL-21 administration (days 14–30) (mean ± SD, *n =* 4 mice). (G) Numbers of lung CD38+CXCR3+CCR6+ and GL7+CD38-B cells from Hif1a^ctrl^ and Hif1a^iKO^ mice at 30 days post-PR8 infection (mean ± SD, *n =* 6 mice). (H) ELISA for PR8-specific IgA from BALF and serum (mean ± SD, *n =* 5*-*6 mice). (I) Numbers of lung CD38+CXCR3+CCR6+ and GL7+CD38-B cells in Hif1a^ctrl^ and Hif1a^iKO^ mice after intratracheal IL-21 treatment (days 14–30) (mean ± SD, *n =* 6 mice). Statistical significance determined by t-tests (A, D, F, I), one-way ANOVA (B, C, E, H), two-way ANOVA (F, G, H) Adjusted p-values: *p<0.05, **p<0.005, ***p<0.0005, ****p<0.0001, ns: not significant. Data representative of two independent experiments.

We next considered if HIF-1α dependent IL-21 might contribute to the maintenance of additional IL-21-sensitive cells in the lung. Pseudobulk analysis indicated that NK cells isolated from influenza infected Hif1a^iKO^ lung at day 30 had decreased expression of *Tcf7*, *Gramd3*, and *Rora*, all shown to be important for the establishment of tissue resident, memory-like cells, while NK cells from control mice were enriched for a signature for ILC1 cells, prominent early producers of IFNγ following viral infection^50–55^ (**Figure S4I**). To assess the lung resident NK cell compartment by flow cytometry, mice were injected intravenously with anti-CD45 shortly before lung harvest. Although the majority of lung NK cells from both Hif1a^ctrl^ and Hif1a^iKO^ mice were in the vasculature (**Figure S4J**), only tissue resident NK cells were decreased in Hif1a^iKO^ mice **(Figure 4E**). In contrast, NK cell numbers in the draining lymph node, spleen and liver were comparable across the strains, indicating that the effects of *Hif1a* deletion are restricted to the lung (**Figure S4K**). A time course analysis in influenza infected wild type mice showed that resident NK cell numbers were elevated at day 9, stably maintained until at least 100 days after infection, and comprised of both conventional (Eomes+) and ILC1 (Eomes-) NK cells **(Figure S4L**). Tissue resident NK cells expressed CXCR6, suggesting possible colocalization with HIF-1α active T cells, and low levels of KLRG1, a marker for terminal NK cell differentiation (**Figure S4M**). Expression of these markers was unaffected in Hif1a^iKO^ mice, suggesting that *Hif1a* deletion affects their recruitment and/or maintenance as opposed to their function **(Figure S4N**). Importantly, administration of exogenous IL-21 to Hif1a^iKO^ mice was sufficient to restore the lung resident NK cell compartment **(Figure 4F**). These data reveal that influenza infection leads to the stable accumulation of lung resident NK cells which depend on IL-21 produced by HIF-1α+ CD4 T cells.

Given the established role of IL-21 in regulating B cell fate, we next examined mucosal B cell responses in Hif1a^iKO^ mice, focusing our analyses on B cells with dual expression of CXCR3 and CCR6, surface markers expressed by antigen specific memory B cells **(Figure S4O**)^37^. Compared to Hif1a^ctrl^ mice, Hif1a^iKO^ mice had lower numbers of lung resident B cells including IgM+, isotype switched (swIg) and IgA+ subsets, as well as GL7+CD38-cells with a germinal center phenotype **(Figure 4G**). B cell subsets in the lung draining lymph node were unaffected, bolstering the idea that late *Hif1a* deletion predominantly affects mucosal as opposed to peripheral immune responses **(Figure S4P)**. Influenza specific IgA titers were also decreased in the BALF of Hif1a^iKO^ mice, contrary to what was observed for IgG **(Figure 4H and S3D**). As shown above, IgA+ cells in wildtype mice were densely localized at the iBALT periphery, similar to the distribution of IL-21+ CD4 T cells from IL-21 VFP reporter mice and HIF-1α-active CD4 T cells from Egln3 reporter mice **(Figures S1B, S4Q and 2D**). Accordingly, intratracheal IL-21 was sufficient to restore these B cell responses in Hif1a^iKO^ mice, highlighting its capacity to rectify broad deficiencies in the lung humoral compartment **(Figure 4I**).

### HIF-1α+ CD4 T cells coordinate mucosal immunity to infectious and tumor challenge

Our data indicate that *Hif1a* deletion in CD4 T cells has wide ranging effects on the influenza induced immune cell network, including lung macrophages, NK cells, and B cells. The coordinated activity of these same cell types was recently reported following heterotypic influenza challenge^31,56^. To probe this further, Hif1a^ctrl^ and Hif1a^iKO^ mice were infected with influenza-X31 followed by tamoxifen administration to delete *Hif1a*, and secondary infection with influenza PR8 **(Figure 5A**). Two days after the challenge, Hif1a^iKO^ mice had decreased oxygen saturation in the blood, a measure of pulmonary function, and increased infectious virus titers **(Figures 5B and 5C**). Both IFNγ expression by lung NK cells and influenza specific IgA titers in the BALF were reduced, underscoring the importance of HIF-1α activity in orchestrating early protective immunity in the mucosa (**Figures S5A and S5B**). To understand if HIF-1α might coordinate a similar network of responses in a distinct context we utilized the KPAR tumor model, a lung adenocarcinoma that elicits TLS and depends on both NK cells and antibodies for tumor control **(Figure 5D)**^57^. Transplantation of KPAR cells into *Egln3*-YFP reporter mice induced the accumulation of lung resident HIF-1α active CD4 T cells, validating the use of this model **(Figure S5C**). Transplantation of KPAR cells into Hif1a^ctrl^ and Hif1a^iKO^ mice followed by tamoxifen treatment led to significantly decreased survival in Hif1a^iKO^ mice (**Figure 5E**). Decreased survival correlated with diminished numbers of FR4+CXCR6+ CD4 T cells, ILC1s and IgA+ B cells in the lung **(Figures 5F-H and 56D**). Similar to influenza infection, AM numbers were decreased in KPAR-bearing Hif1a^iKO^ mice **(Figure 5I**). Hif1a^iKO^ mice also harbored more MHC-II-low myeloid cells which have been associated with compromised immunity in the tumor microenvironment **(Figure 5J**)^58^. Remarkably, survival in Hif1a^iKO^ mice could be prolonged by the exogenous administration of IL-21 **(Figure 5K**).

**Figure 5.**
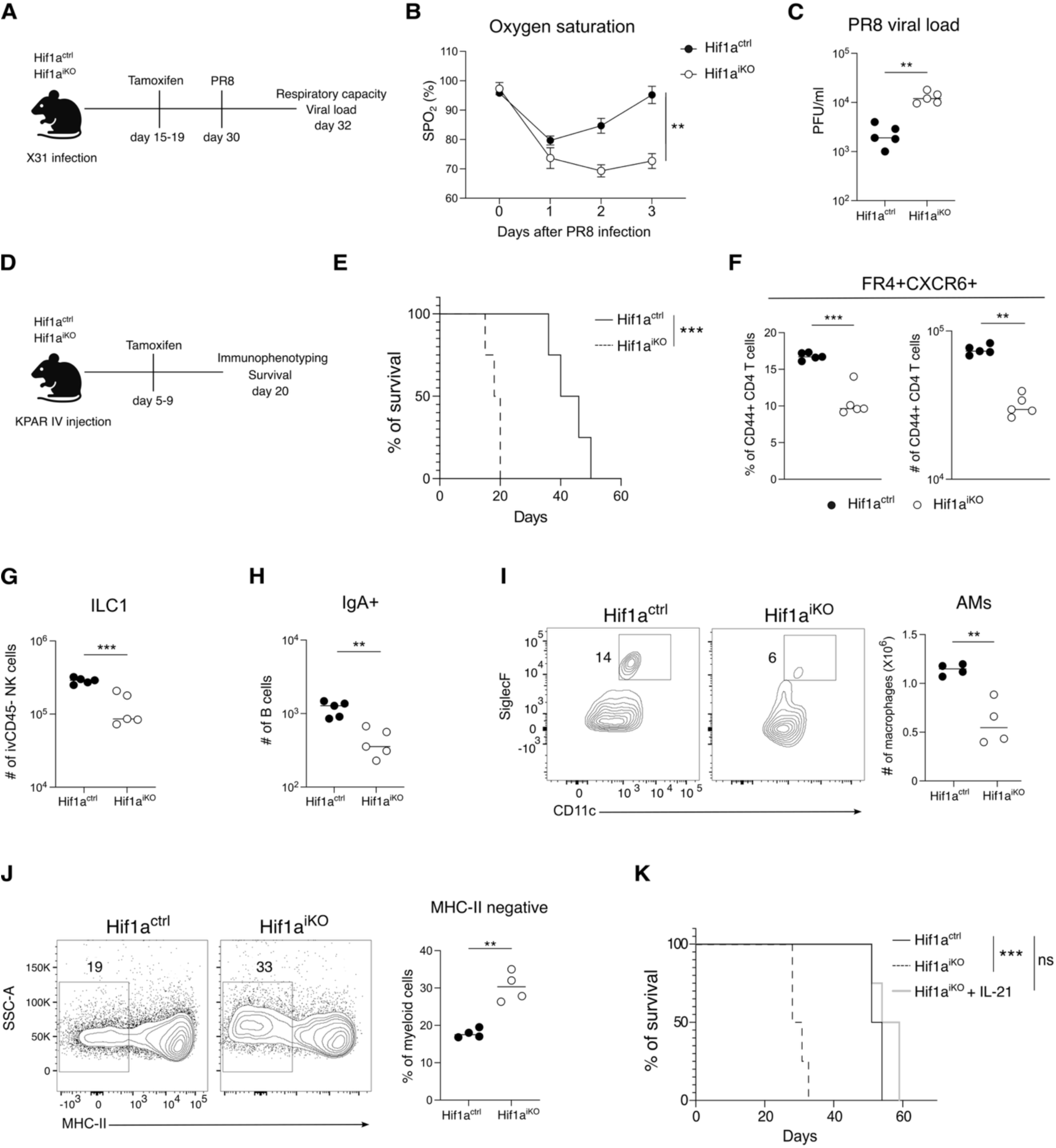
HIF-1α+ CD4 T cells coordinate mucosal immunity to infectious challenge and tumor. (A) Experimental design for the analyses shown in (B-C). (B) Measurement of oxygen saturation (SpO_2_) after challenge (mean ± SD, *n =* 5 mice). (C) Viral plaque assay with MDK2 cell line for Hif1a^ctrl^ and Hif1a^iKO^ mice after challenge (mean ± SD, *n =* 5 mice). Experimental layout for the data in (E–J). (E) Survival curve of Hif1a^ctrl^ and Hif1a^iKO^ mice injected with KPAR cells (mean ± SD, *n =* 5 mice). (F) Frequency and number of FR4+CXCR6+ CD4 T cells (mean ± SD, *n =* 5 mice). (G) Numbers of lung ILC1 cells and (H) IgA+ resident B cells (mean ± SD, *n =* 5 mice). (I) Representative plot and numbers of AMs (mean ± SD, *n =* 4 mice). (J) Frequency of MHC-II negative of lung myeloid cells from Hif1a^ctrl^ and Hif1a^iKO^ mice (mean ± SD, *n =* 4 mice). (K) Survival curve of Hif1a^ctrl^ and Hif1a^iKO^ mice following intratracheal IL-21 treatment (days 14–30) (mean ± SD, *n =* 4 mice). Statistical (mean ± SD) significance determined by t-test Aggregate comparison AUC (B), Mantel-Cox test (E) and t-tests (C, F-J). Adjusted p-values: **p<0.005, ***p<0.0005, ns: not significant. Data representative of two independent experiments.

## DISCUSSION

The tissue microenvironment is composed of a variety of resident immune cells working in concert to maintain tissue homeostasis. During infection, T cells in the tissue and TLS may re-encounter antigen, respond to inflammatory cytokines, engage with new cell types, and experience fluctuations in nutrient and oxygen availability. T cell plasticity - a cornerstone of the immune response - allows T cells to sense and adapt to these dynamic, tissue derived signals. This plasticity is facilitated by the flexible expression of transcription factors that fine tune T cell function. However, genetic models to conditionally delete transcription factors in tissue T cells are lacking. In this study, inducible deletion of *Hif1a* in CD4 T cells at the height of its activity in the lung restricts the effect to the tissue, revealing HIF-1α’s role as a nexus of tissue T cell plasticity.

During influenza, HIF-1α is most highly expressed by lung TRH cells, which share phenotypic and functional characteristics with lymphoid Tfh cells, and which exhibit the flexibility to differentiate into both TRH and TRM1 cells following adoptive transfer and challenge^4,5^. We demonstrate that HIF-1α+ CD4 T cells contribute to TRH plasticity via their production of IL-21, which leads to CXCR6 upregulation and seeding of tissue resident cells at the iBALT perimeter. An open question is whether IL-21 produced by lung TRH cells acts in an autocrine or paracrine manner to upregulate CXCR6. A recent study showed that IL-21 produced by lymphoid Tfh cells can act in *trans*, indicating that both scenarios are possible^59^.

Future efforts should focus on elucidating the signals that drive *Hif1a* upregulation in lung CD4 T cells. Although our data suggest that HIF-1α+ CD4 T cells arise locally from lung TLS, TLS themselves persist long after HIF-1α activity has waned, indicating that while TLS are necessary, they are not sufficient for the maintenance of HIF-1α+ CD4 T cells^60^. Importantly, HIF-1α in T cells can be elicited by a variety of inputs including hypoxia, T cell receptor signaling and TGFβ, all of which were detected at the TLS border by spatial transcriptomics^18,21,22,61–63^. It is possible that the final differentiation outcome of Hif1α activity and IL-21 sensing depends on additional, spatially compartmentalized signals that stabilize one fate over another^64,65^. A parallel can be drawn with B cells, where IL-21 regulates both Bcl6 and its antagonist Blimp1, balancing high affinity selection in germinal centers with plasma cell differentiation^66–68^. For tissue T cells, IL-21 and cognate antigen presentation by B cells in the TLS core might reinforce TRH cell identity, while interactions with CXCL16 expressing antigen presenting cells outside of TLS may favor Blimp1, required for tissue residency and a predicted regulator of TRM1 cell identity^69,70,4^. Alveolar macrophages and epithelial cells - both sources of CXCL16 - may contribute to this specially organized regulation of tissue resident T cells^45,71,72^. It is also worth noting that in the gut mucosa, deletion of *Hif1a* in T cells has pleiotropic effects across Th17, Tfh cells and regulatory T cell subsets, all of which are capable of interconverting^73,12,74,75,10^. It is tempting to speculate that HIF-1α plays a broader role in modulating T cell plasticity. Given that Bcl6 can antagonize certain aspects of HIF-1α activity, co-expression of these transcription factors may mark a poised state, enabling rapid engagement of a HIF-1α-driven transcriptional program upon downregulation of Bcl6^76^. An additional layer of control provided by HIF-1α may account for the functional redundancy among Th subsets when an individual lineage determining transcription factor is deleted^77–79^.

Our results underscore a role for HIF-1α+ CD4 T cells in establishing a mucosal immune cell network including replenishment of the alveolar macrophage compartment, accumulation of lung resident NK cells, and maintenance of humoral immunity in the respiratory tract. These diverse responses converge on a shared requirement for IL-21 provided by Hif1α+ CD4 T cells. A similar network has recently been described in which influenza specific memory B cells, persisting in alveolar as opposed to iBALT niches, require both alveolar macrophages and NK cells in order to migrate to infected foci after challenge^31,56^. Our data suggest that HIF-1α+ CD4 T cells may coordinate an analogous IL-21 dependent immune cell network within the tumor microenvironment. Notably, these data are aligned with the correlation between the intratumoral expansion of IL-21+ CD4 T cells and successful outcomes to immune checkpoint blockade^80,81^.

In summary, HIF-1α activity is central to CD4 T cell plasticity in the tissue and facilitates IL-21 dependent control over a multifaceted immune response in the lung. TLS and surrounding microenvironmental cues emerge as key sites for the differentiation of HIF-1α+ CD4 T cells, although downstream effector functions are likely shaped by additional signals encountered outside the TLS core. While these findings highlight a tissue-specific role for HIF-1α, they do not exclude its involvement in early peripheral lymphoid responses, and may help resolve some apparently contradictory effects of HIF-1α deletion on Th cells.

## RESOURCE AVAILABILITY

### LEAD CONTACT

Further information and requests for resources and reagents should be directed to and will be fulfilled by the lead contact, Carolyn King.

## DATA AND CODE AVAILABILITY

Single-cell RNA-seq data have been deposited at GEO: accession number and are publicly available as of the date of publication. Microscopy data reported in this paper will be shared by the lead contact upon request. Any additional information required to reanalyze the data reported in this paper is available from the lead contact upon request.

## ACKNOWLEDGEMENTS

We thank G. Auray for cell sorting, D. Calabrese for histology support, core facilities (microscopy and bioinformatics) as well as animal caregivers at the Department of Biomedicine, and members of the King lab for helpful discussion. This work was supported by the Swiss National Science Foundation, grant 197683 to CGK, National Institutes of Health grants AI154598, AI147394, HL170961 and AI176171 to JS and NIH grant R00 HL157555 to ATS.

## AUTHOR CONTRIBUTIONS

Conceptualization: JDL, CGK; Methodology: JDL, DS, CGK; Formal analysis: JDL, EB, DS, CGK; Investigation: JDL, NS, JS, ME, LL, ME(2), LL, CED, MK, ATS, DS; Resources: AG, JS; Visualization: JDL, DS, CGK; Funding acquisition: CGK; Supervision: DS, CGK; Writing: JDL, DS, CGK.

## SUPPLEMENTAL INFORMATION

Document S1. Figures S1-S5.

## STAR * METHODS

### KEY RESOURCES TABLE

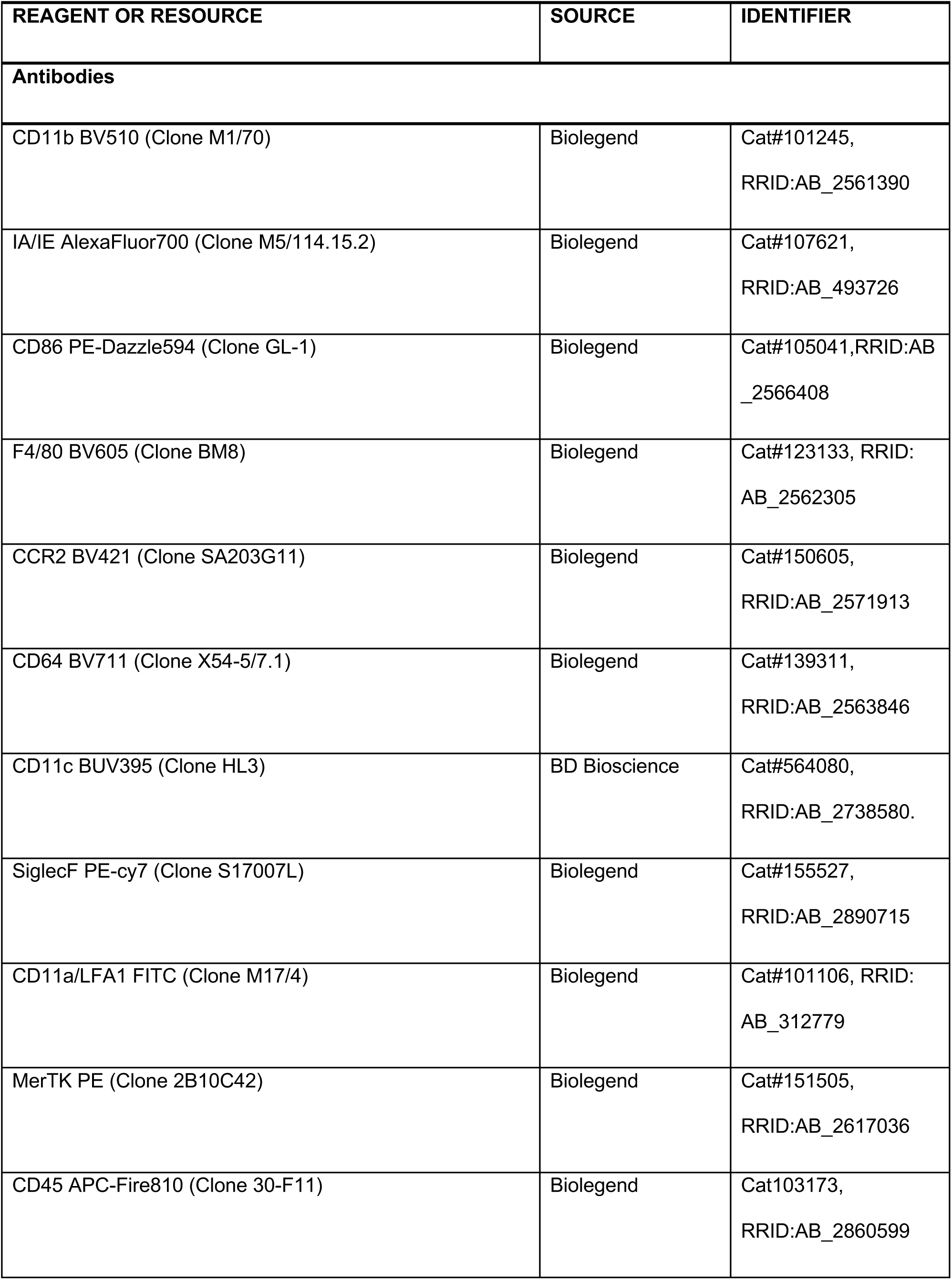
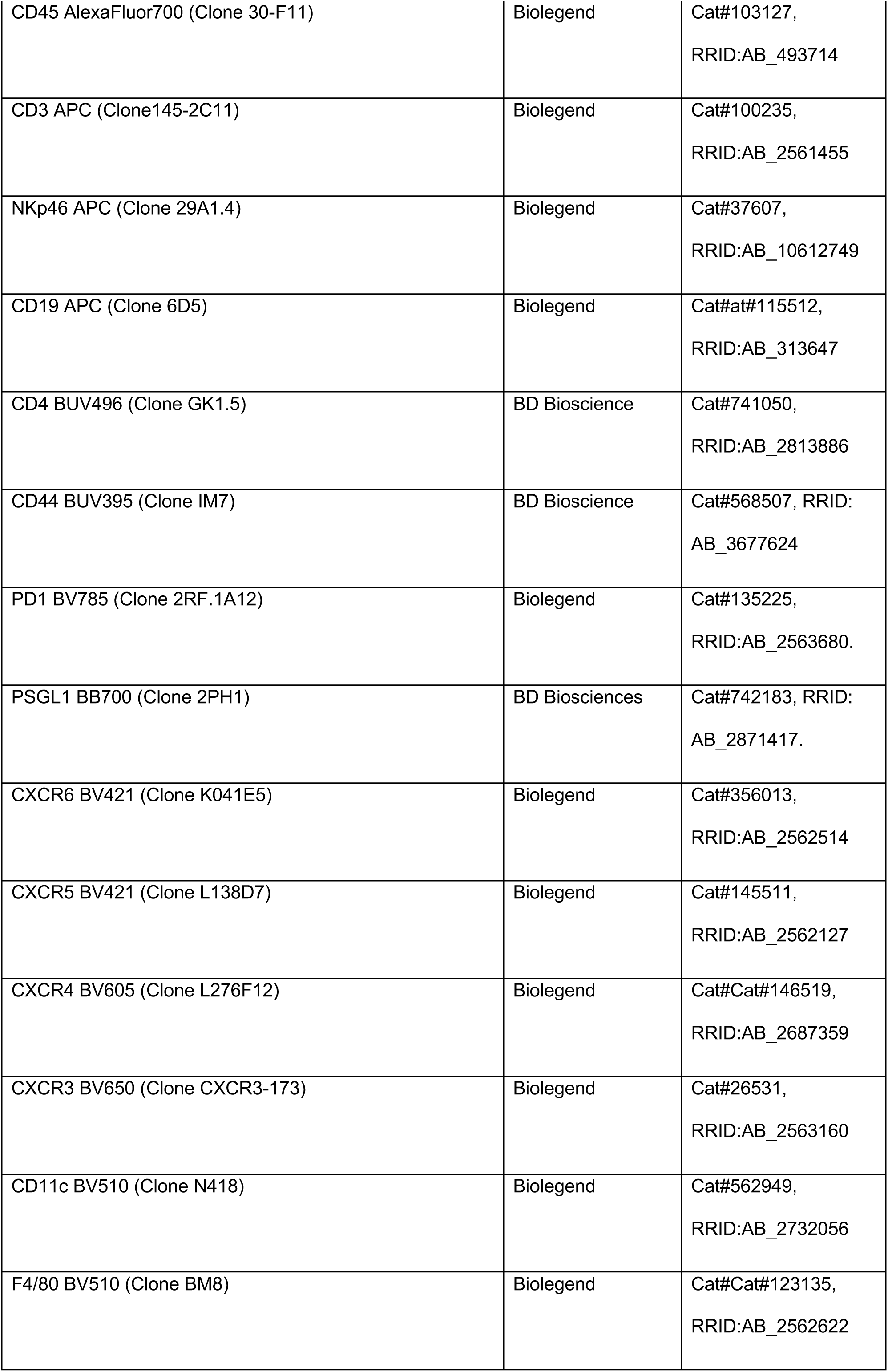
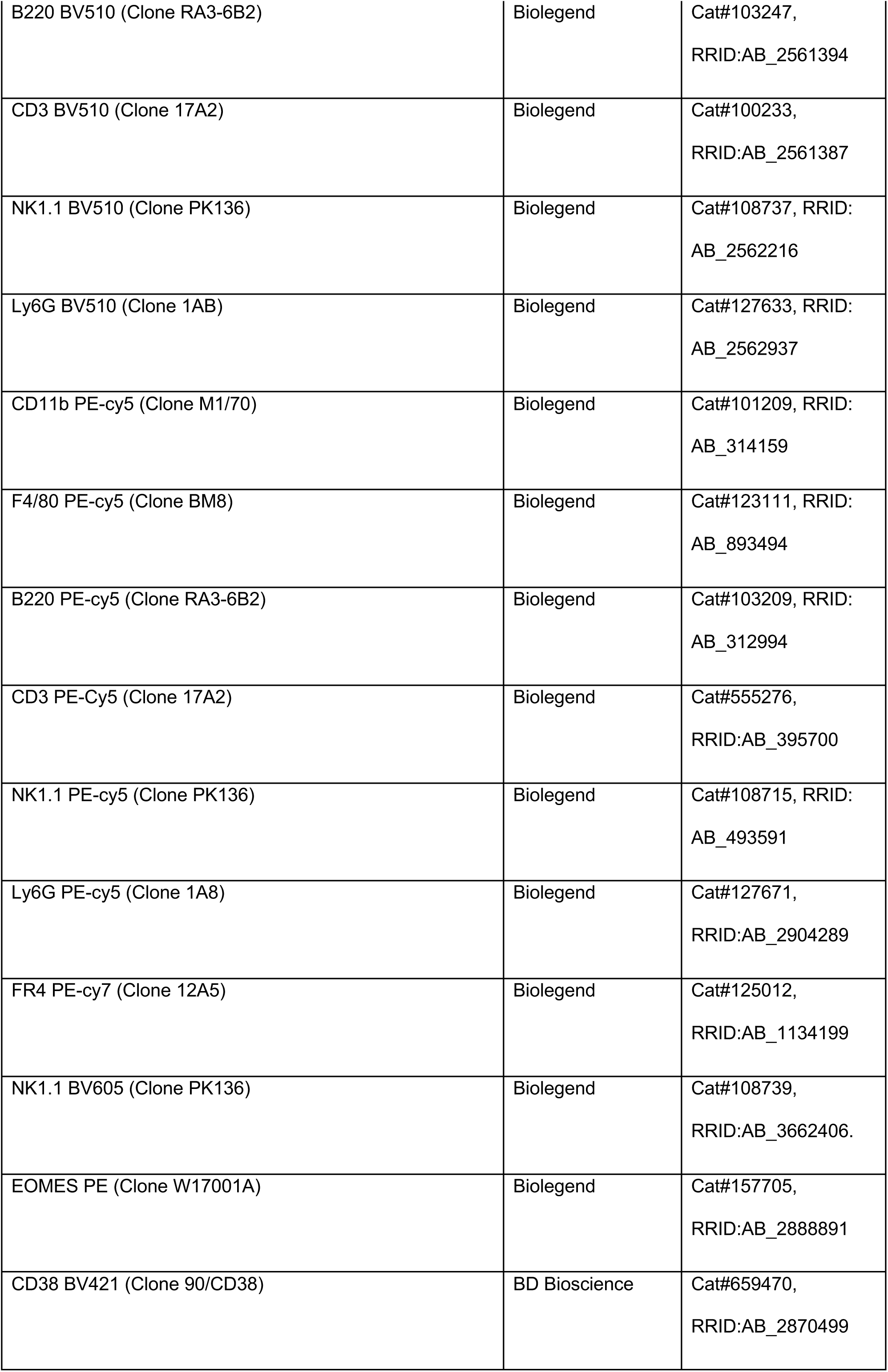
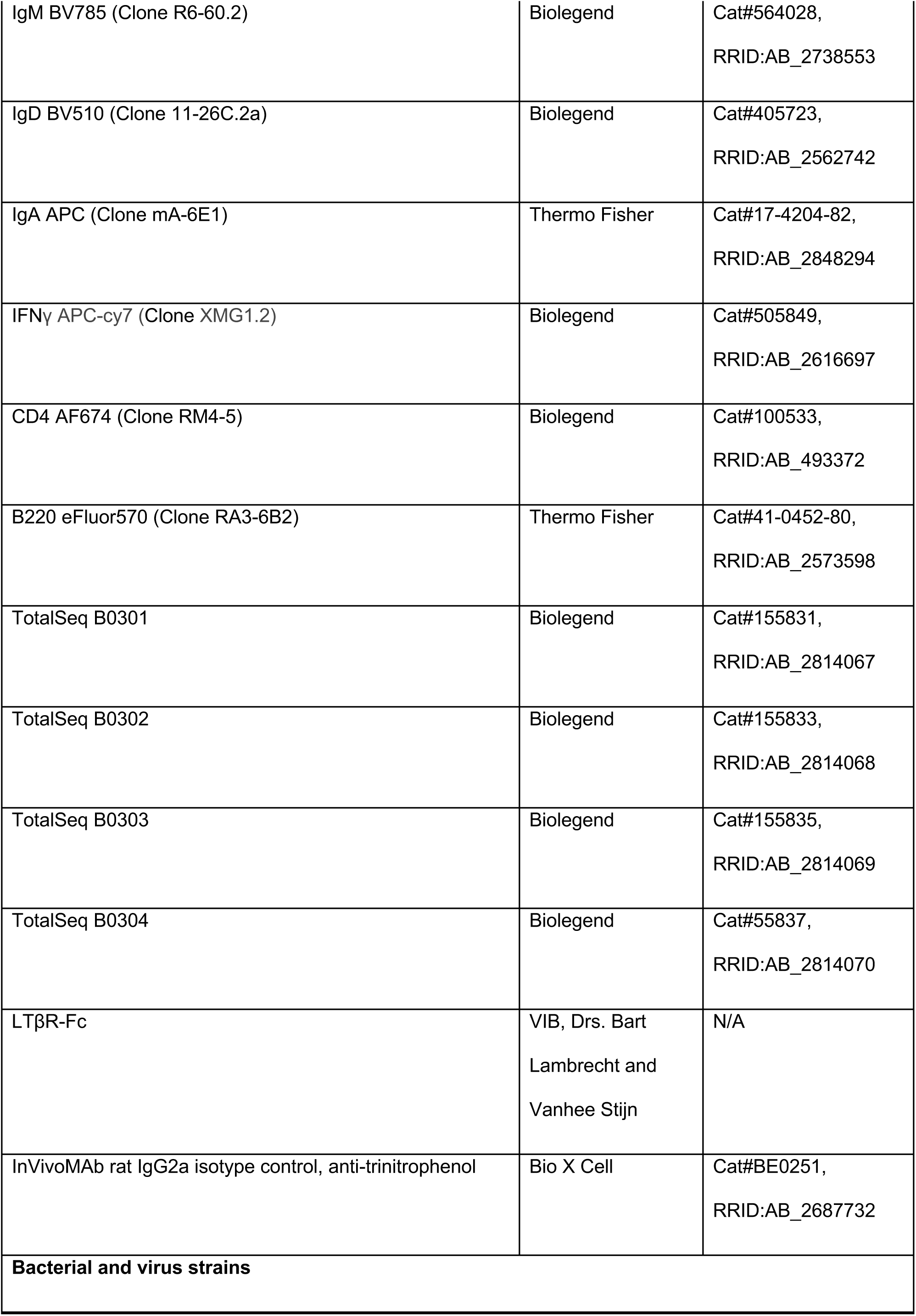
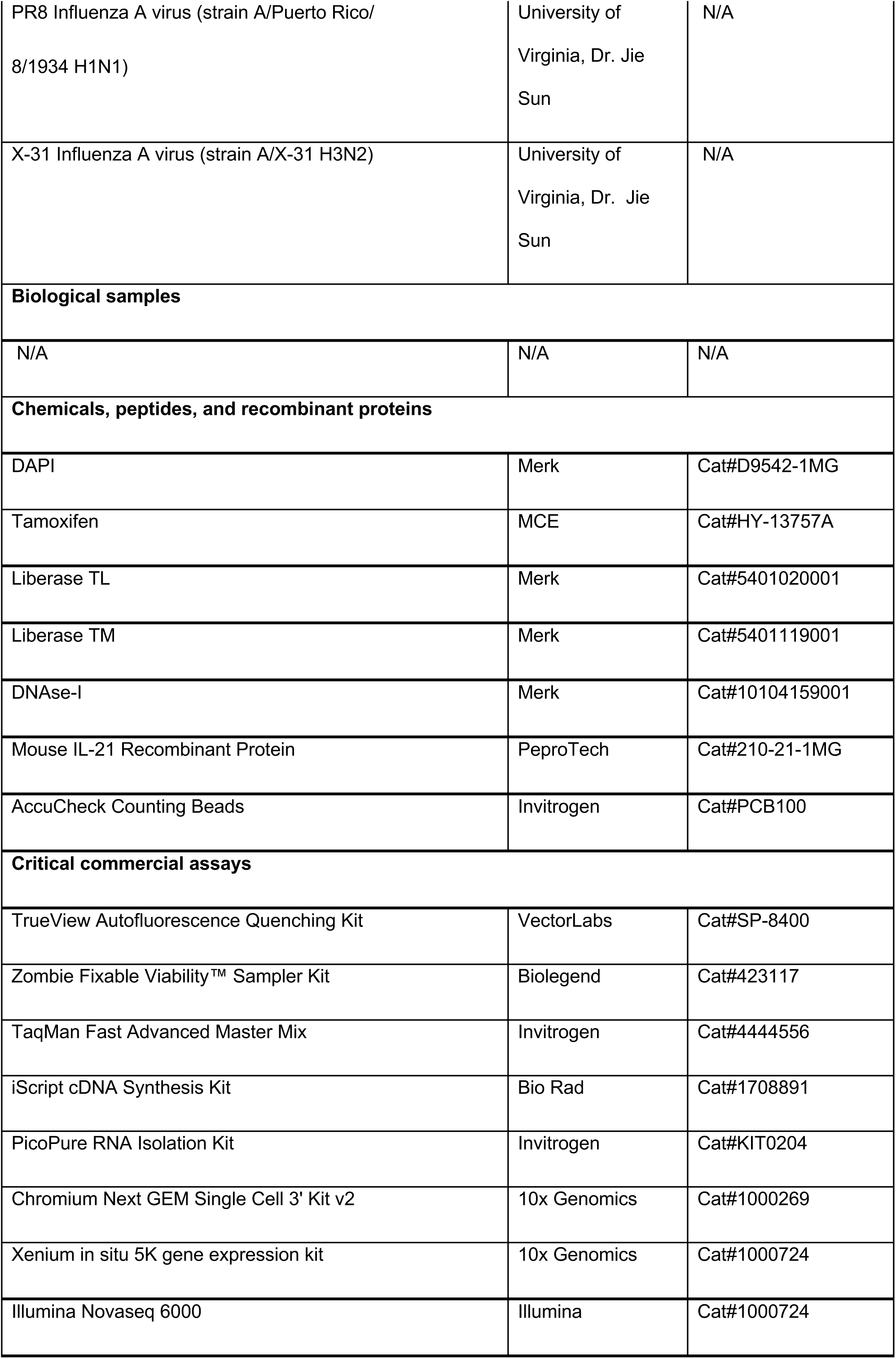
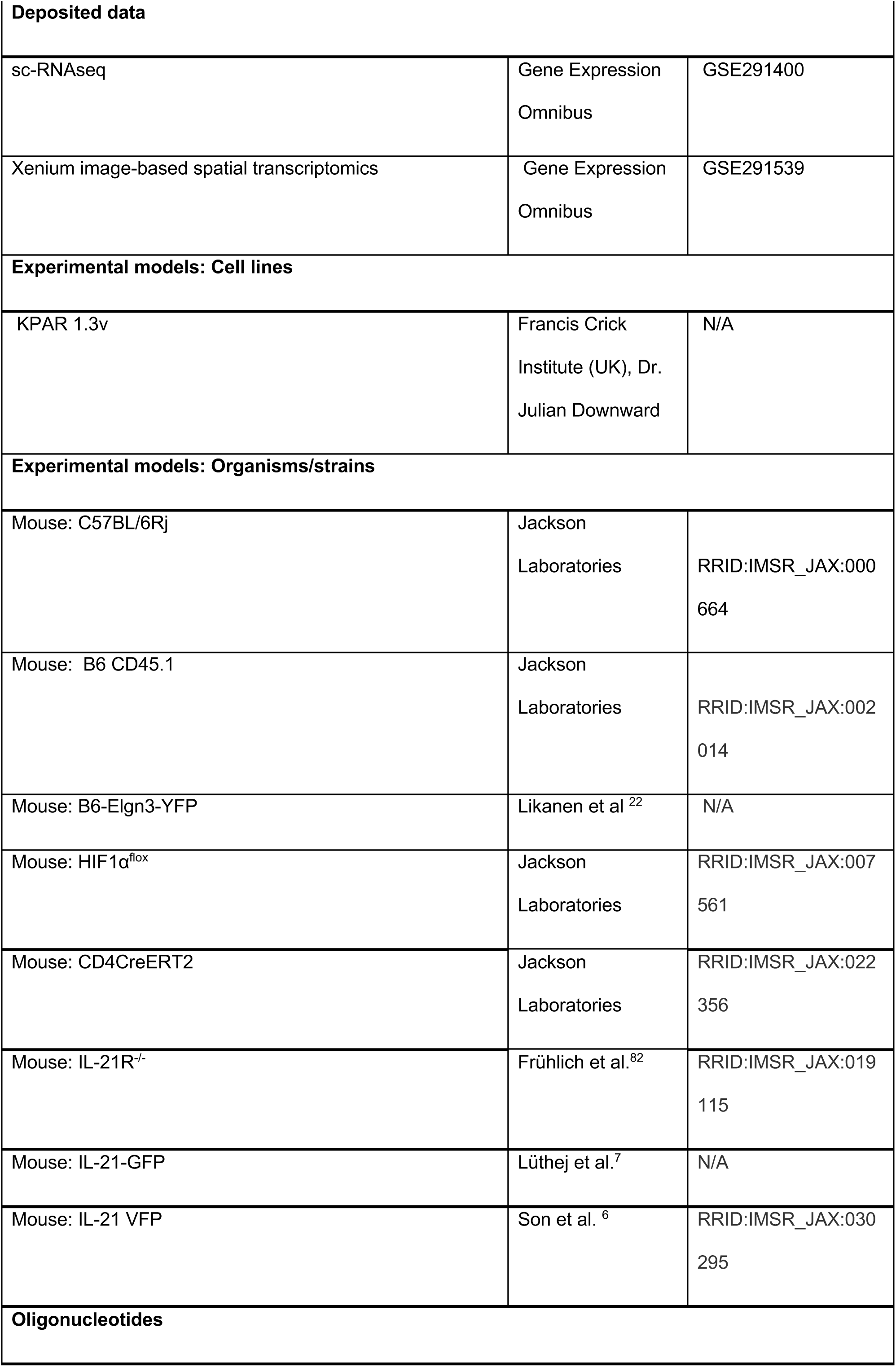
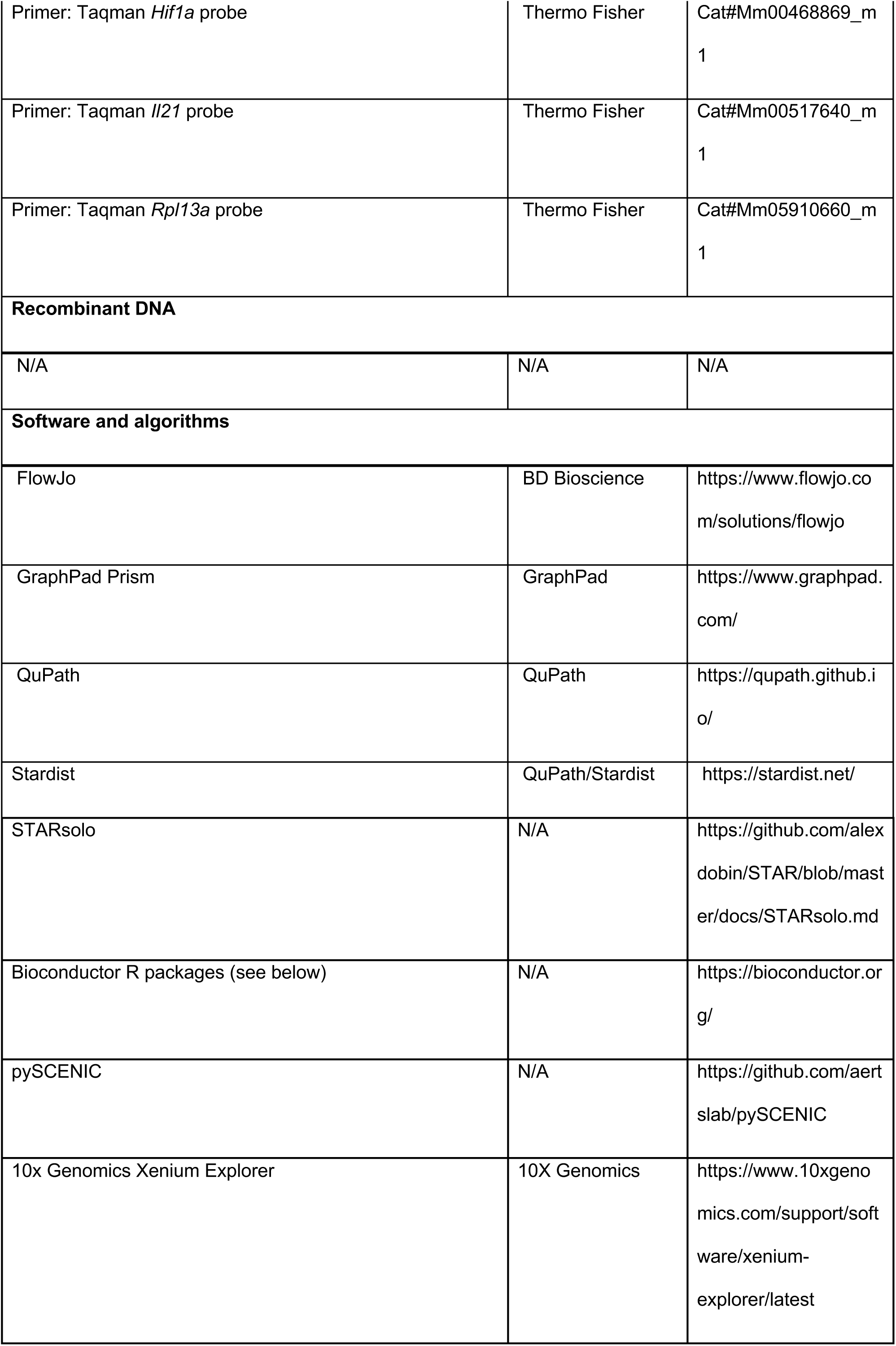
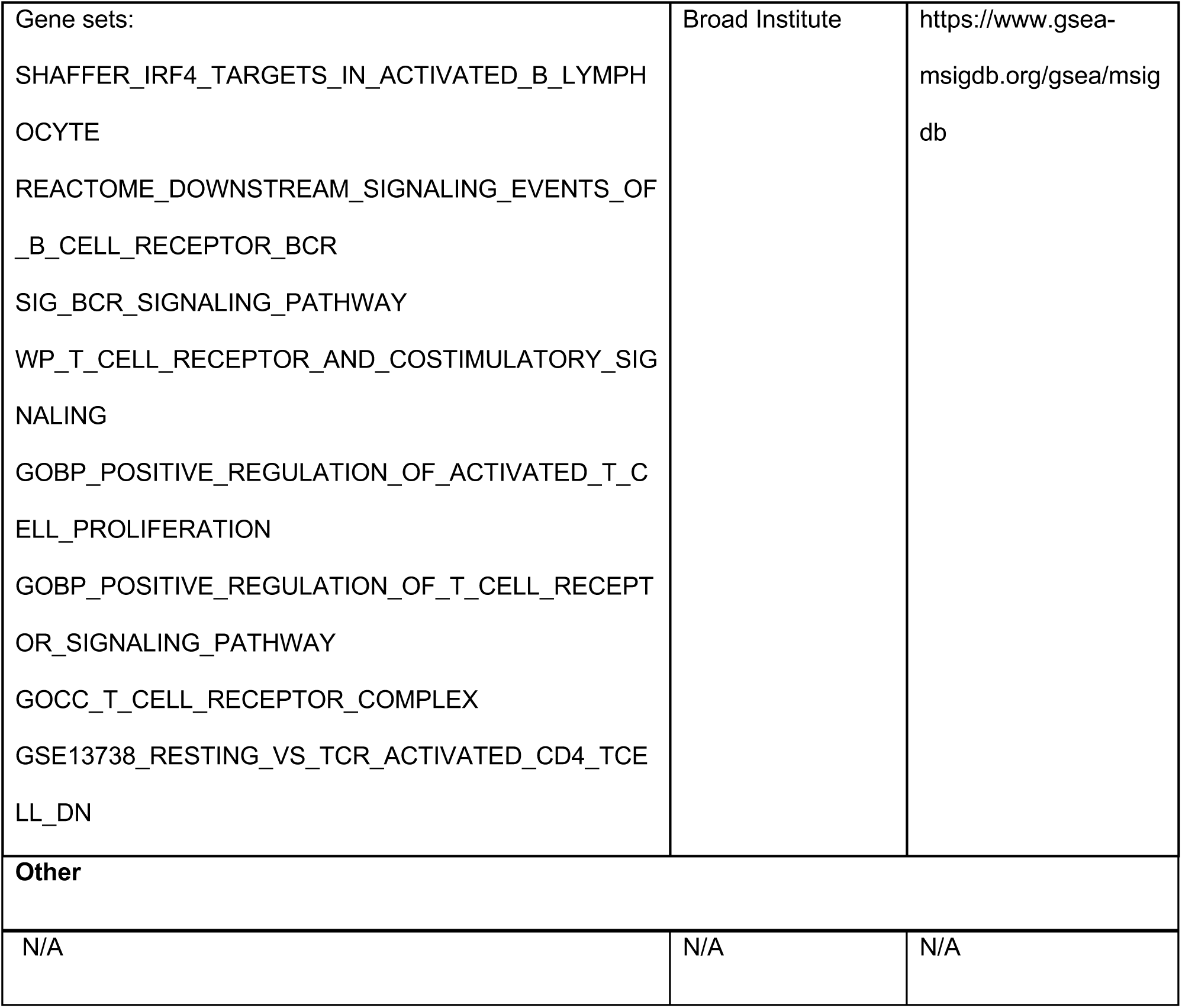

## EXPERIMENTAL MODEL AND STUDY PARTICIPANT DETAILS

### Mice

C57BL/6Rj (CD45.2) were purchased from Janvier Labs. *Egln3-YFP* mice, B6-Elgn3<tm1YFPAgo>, were generated by A. Goldrath^22^. *Hif1a*^flox/flox^ mice^83^ were crossed with *CD4*^ERT2^ x *Rosa*-26 mice^84^. IL-21R^-/-^ mice, <B6.129-*Il21rtm1Kopf*/J>, were generated by M.Kopf^82^. IL-21-GFP mice were generated by S. Nutt^7^. Mice in each experiment were 6-8 weeks old, same-sex littermates, maintained and bred in the specific pathogen– free animal facility at the University of Basel. All animal experiments were performed in accordance with local and Swiss federal guidelines.

### Influenza infection

Influenza A/PR8/34 (hereafter PR8) and X31 were produced by J. Sun, University of Virginia. PR8 was used at 500 PFU/TCID50 and X31 at 800 PFU/TCID50. Mice were anesthetized with vaporized isoflurane and intratracheally infected intratracheally with virus diluted in 50 μL of PBS.

### Bone Marrow Chimera

To produce mixed chimeras, recipient B6-CD45.1 mice were lethally irradiated with two doses of 450 cGy given 4 hours apart. After irradiation, recipients were reconstituted by intravenous injection of hematopoietic cells collected from femurs and tibiae of donor mice (50% B6-CD45.1 and 50% IL-21R^-/-^), as described before^85^. Mice were used for experiments 6–8 weeks after irradiation.

### KPAR tumor

KPAR cells were kindly provided by Julian Downward, Francis Crick Institute, London ^57,86^. 2.5X10⁵ KPAR cells (maintained at a maximum of 5 passages) were intravenously injected into the tail vein of each mouse.

## METHOD DETAILS

### In vivo treatments

Tamoxifen (Sigma-Aldrich) dissolved in corn oil (25 mg/ml) and administered by oral gavage (3.75 mg/mouse) at indicated time points. Anti-CD45 AF700 (Biolegend) was intravenously injected (3 μg/mouse) just prior to sacrifice and organ harvest. Recombinant mouse IL-21 protein (PeproTech) was intratracheally administered every other day (10 μg/treatment/mouse) for the indicated time points. LTβR-Fc was generated and kindly provided by Bart Lambrecht and StijnVanhee. Mice were intraperitoneally injected with 100ug/mouse for the indicated time points.

### Tissue preparation

Lungs were harvested and diced into gentleMACS C Tubes (Miltenyi Biotec) and washed down with 3 ml of media [RPMI, 10 mM aminoguanidine hydrochloride (Sigma-Aldrich), 10 mM Hepes, penicillin-streptomycin-glutamine (100×, Gibco), 2-mercaptoethanol (50 μM, Gibco)], prewarmed in a 37°C water bath. Digestion mix containing liberase TL (for lymphoid cells) or liberase TM (for myeloid cells) (33.3 μg/ml, Merk) and deoxyribonuclease I (58 μg/ml; Applichem) was added. Lungs were dissociated on a gentleMACS Dissociator (Miltenyi Biotec) and mashed through 70-μm MACS SmartStrainers as previously described (Miltenyi Biotec)^4^. Mouse mediastinal LNs and spleens were harvested and mashed through 100-μm Corning nylon cell strainers. For scRNA-seq experiments, mediastinal LNs were additionally digested with liberase TL. Mouse livers were perfused with PBS and filtered through 100-μm Corning nylon cell strainers. Lymphocytes from the liver were further purified on a 70%/40% Percoll gradient (Fisher Scientific).

### Tetramer and flow cytometry staining

Ι−Ab NP311-325 (QVYSLIRPNENPAHK) allophycocyanin (APC-) conjugated was obtained from the National Institutes of Health (NIH) tetramer core facility. Single-cell suspensions were tetramer-stained, enriched, and counted as previously described(*61*). For all fluorochrome-conjugated antibody dilutions, FACS buffer (PBS, 2% FCS, and 0.1% sodium azide) containing Fc block (Clone 2.4G2 InVivoMAb anti-mouse CD16/CD32; BioXCell, #BE0307) was used. Live-dead stain (provided by Zombie Fixable Viability™ Sampler Kit; BioLegend) was added to the antibody mix and stained at 4°C for 30 min. FoxP3 Transcription Factor Staining Buffer Set (eBioscience) was used for intracellular staining at room temperature (RT) 30-60 minutes. Flow cytometric analysis was performed on BD LSR Fortessa and Cytek Aurora. Data was analyzed using FlowJo X software (BD) and SpectraFlow (Cytek). Absolute cell numbers were based on AccuCheck Counting Beads (Invitrogen) manufacturer.

### Confocal microscopy

Lungs were inflated with a 1:1 mixture of Tissue-Tek® OCT. Compound (Sakura) and PBS containing 1% methanol-free PFA (Thermo Fisher Scientific) and fixed with 4% PFA for 24 hours. Lungs were washed in PBS, incubated in a 30% sucrose solution for 36 hours and embedded in OCT. For immunostaining of lung sections from *Egln3-YFP* mice, tissues were sectioned at 10-30 μm on a cryostat and incubated with 5% blocking buffer (Jackson ImmunoResearch) for 2 hours. Slides were incubated with indicated antibodies overnight at 4°C followed by washing with PBS, DAPI staining (2 μg/mL, Merck) and treatment with TrueView Autofluorescence Quenching Kit (Vector) to reduce autofluorescence. Sections were mounted using SlowFade Diamond Antifade Mountant (Thermo Fisher) to preserve *Egln3-YFP* fluorescence. Z-stack images (10μM to 30μM, step size of 0.9μM) were acquired on a Nikon ECLIPSE Ti2 inverted microscope, equipped with a X-light V3 spinning disk confocal module and a 40X Apo Plan lambda, NA=0.95 objective.

### Image quantification

Multichannel fluorescence images were analyzed with QuPath (ver. 0.4.3)(*62*, *63*). Cells were segmented based on nuclear DAPI staining using a pre-trained deep learning model cyto3(*64*) from cellpose, with parameters: pixel size: 0.33; object diameter: 30, cellExpansion: 2.0. Positivity of cells (Biolegend CD4 AF647 and *Egln3*-YFP) was determined based on the mean intensity measured within the cell (ROI). The B cell cluster was identified using a pixel classifier based on B220 staining. A training image was composed of a representative set of 6 images which was annotated by a user to identify positive and negative B220 (Thermo Fisher, eFluor-570) signals. ANN_MLP classifier was trained with the following parameters: resolution: 2.66µm; filters: Gaussian and Laplacian; scales: 1 and 4. Finally, the signed distance of each cell to the B cell cluster annotation was calculated using the detectionToAnnotationDistancesSigned function, and individual cell distance measurements were extracted in a csv file.

### Real-time PCR

RNA was extracted from sorted NP+ TRM1, Egln3-TRH cells, and Egln3+ TRH cells using PicoPure^TM^ RNA Isolation Kit (Thermo Fisher). RNA was reverse transcribed into cDNA using iScript cDNA (Synthesis Kit, Bio-Rad). Taqman Fast Advanced probes (Applied Biosystems) for *Il21* (#Mm00517640_m1), *Hif1a* (#Mm00468869_m1) and *Rpl13a* (#Mm05910660_g1). were used for this assay. For data presentation, relative expression was calculated and normalized to *Rpl13a.* Fold change values were calculated using the 2^-ΔΔCT^ method.

### Blood oxygen saturation (SpO_2_)

Blood SpO_2_ was measured using a MouseOx™ pulse oximeter (Starr Life Sciences). Anesthetized mice were placed on a heating pad to maintain body temperature and an oximeter paw sensor was secured to the hind limb using medical adhesive tape. During the recording, SpO_2_ measurements were taken 3 times, each measurement lasting approximately 10 minutes.

### Viral titers during influenza challenge

Mice were infected with the X31 influenza strain for 30 days followed by secondary infection with PR8 influenza. Viral titers in the BALF were measured 36 hours after challenge as previously described^91^.

### ELISA

The coating method for plates, antibody concentrations used, and the steps for detecting PR8-specific IgG and IgA were performed as previously described^92^.

### Single cell RNA sequencing

Following influenza infection, control (CD4^Cre-ERT2^ROSA^ki/+^) and Hif1a^flox/flox^CD4^Cre-ERT2^ mice were treated with tamoxifen to induce gene deletion in CD4 T cells. Individual mice were hashtagged using TotalSeq-B anti-mouse hashtag antibodies (B0301-B0304, BioLegend), and CD45+ cells were sorted from lungs at d45 p.i. and submitted for library preparation. A total of 10,000 target cells per sample were loaded. Single-cell capture and cDNA library preparation was performed according to manufacturer’s instructions using 10x Genomics Chromium Next GEM Single Cell 3’ Kit v2 reagents using dual indexing. Sequencing was carried out on an Illumina Novaseq 6000 at the Genomics Facility Basel, ETH Zurich.

Sequencing reads were mapped to the mouse transcriptome (mm39) using STARsolo for both gene expression and antibody-bound hashtag oligos using cell barcode whitelist from cellranger-7.0.0. Unless otherwise noted, analysis steps were performed using Bioconductor packages: SingleCellExperiment, scater, scran, scuttle, bluster, DropletUtils, SingleR, edgeR, limma^93–100^. Visualizations with ggplot^101^. Demultiplexing hashtag oligos and doublet detection were done using DropletUtils. Cells were further filtered for library size, library complexity and mitochondrial content using adaptive thresholding provided by the R package scuttle’s perCellQCFilters function with default arguments. Normalization was performed using the R package scran’s deconvolution method. Technical noise within gene expression was modeled using scran, and biologically relevant highly variable genes (HVG) were used after separating the technical from biological variance. Principal component analysis (PCA) was run on the normalized data using the top 2000 most variable genes by biological variance, and the PCA was denoised to account for the modeled technical variation. Cells were clustered using the bluster R package, either hierarchically using Ward’s method or with the graph-based leiden method. Cell annotation was performed using the SingleR package with references from the celldex package followed by manual curation based on cluster-identifying genes enabled the identification of NK/ILC1 cells. Secondary analysis of published datasets was downloaded from Gene Expression Omnibus (GEO). Single-cell regulatory network inference and clustering (SCENIC) was performed on data from GEO dataset GSE146626 using version 0.9.7 and the published workflow on the pySCENIC github repository^102,103^. GSEA between conditions was performed using camera from the limma R package on standard gene set categories (MSigDB) as well as sets curated from relevant publications.

### Spatial transcriptomics

Lungs were fixed with 4% methanol-free PFA for 24 hours at 4°C, washed with PBS and paraffinized using Tissue Processing Center (TPC 15 Trio, Medite) and embedded in paraffin using TES station machine (Tissue Embedding Station, Medite). FFPE blocks were kept at 4°C and 5μm sections and placement of sections were performed according to the protocol CG000578 (10X Genomics), handled by the histology facility of Basel. Xenium Prime 5K processing and acquisition (with cell segmentation) was performed by Functional Genomics Center Zurich, following the user guides GC000760, GC000749 and GC00582 using Xenium in situ 5K gene expression kit (10 Genomics).

Probability-based segmentation was performed using Proseg as an alternative to 10x Genomics’ multi-modal segmentation^104^. Analysis steps such as finding HVGs, PCA, cell annotation and clustering were done with bioconductor R packages as in sc-RNAseq processing. Identification of iBALT: normalized expression of *Cd79b* was summed for the 32 nearest neighbors of each cell. Every cell with a log10 neighbor expression higher than 1.05 was deemed an iBALT cell (threshold determined with manual comparison to H&E image). The general idea was to find areas not only where an individual cell has high *Cd79b* expression, but also where it is surrounded by "enough" other cells that also have high *Cd79b* expression. iBALT cells were then clustered hierarchically based on their X and Y coordinates to define discrete iBALT structures across the section. Cells within 300 µm of iBALT cells that were themselves not also iBALT cells were considered to be in the iBALT periphery, and the distance to nearest iBALT edge was recorded for iBALT and iBALT periphery cells (with iBALT and iBALT periphery cells receiving negative and positive distance values respectively). Gene set scores calculated using the UCell R package.

## QUANTIFICATION AND STATISTICAL ANALYSIS

Prism software (GraphPad 10. 4.1) was used for all statistical analyses. All statistical tests used are two-tailed unpaired t-tests, one/two-way ANOVA, Aggregate comparison (AUC) and Mantel-Cox test, which are assigned to each individual figure panel legend. Statistical significances are indicated by \**p*<0.05, \*\**p*<0.005, \*\*\**p*<0.0005, \*\*\*\**p*<0.0001, ns, not significant (*p* > 0.05). In summary graphs, points indicate individual samples, and horizontal lines are mean unless otherwise noted. All error bars are standard deviation.

---

**Figure S1.**
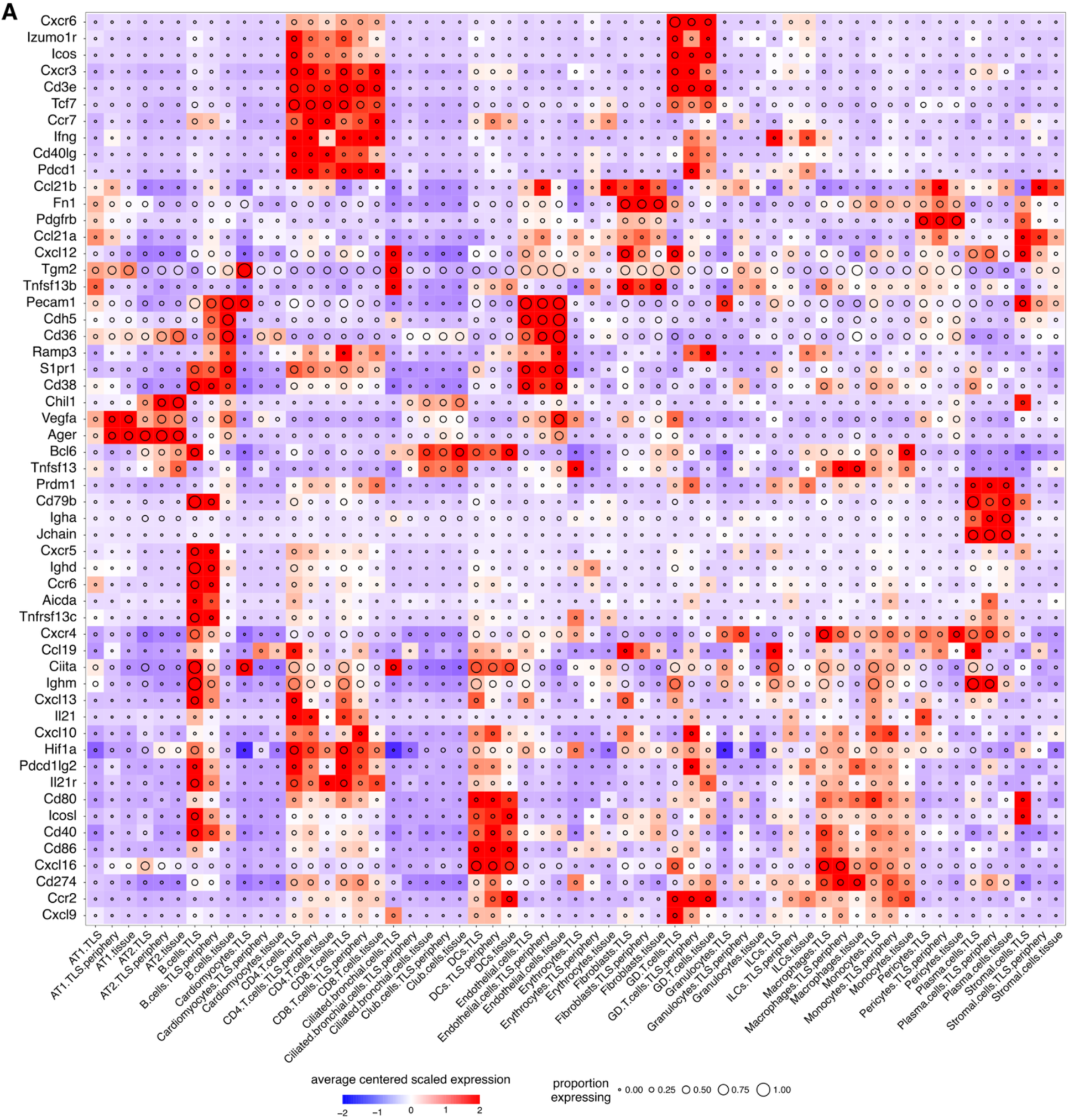
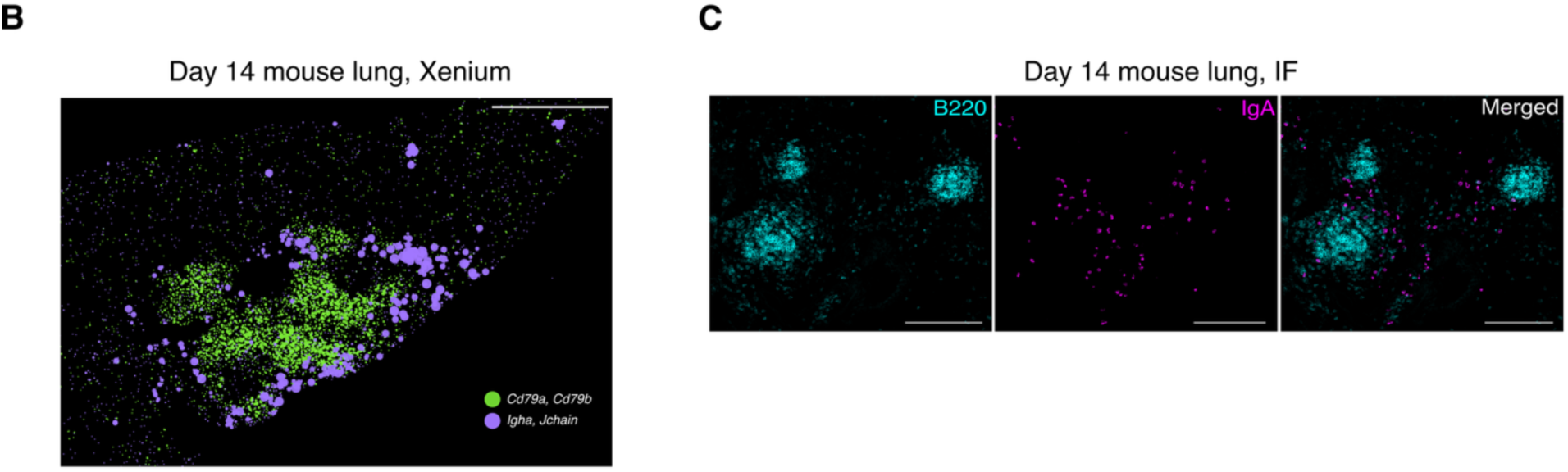
Spatially structured immune cell interactions in lung TLS. (A) Centered, scaled normalized expression of Xenium data of mouse lung day 14 post-influenza. Expression averaged per cell type and tissue location (*n =* 1 mouse). (B) Representative iBALT region from day 14 lung as displayed in Xenium Explorer. Bars, 500µm. (C) Immunofluorescence (IF) showing inducible iBALTs (cyan) containing IgA⁺ cells (magenta) 14 days after influenza infection. Bars, 200µm (*n =* 3 mice).

**Figure S2.**
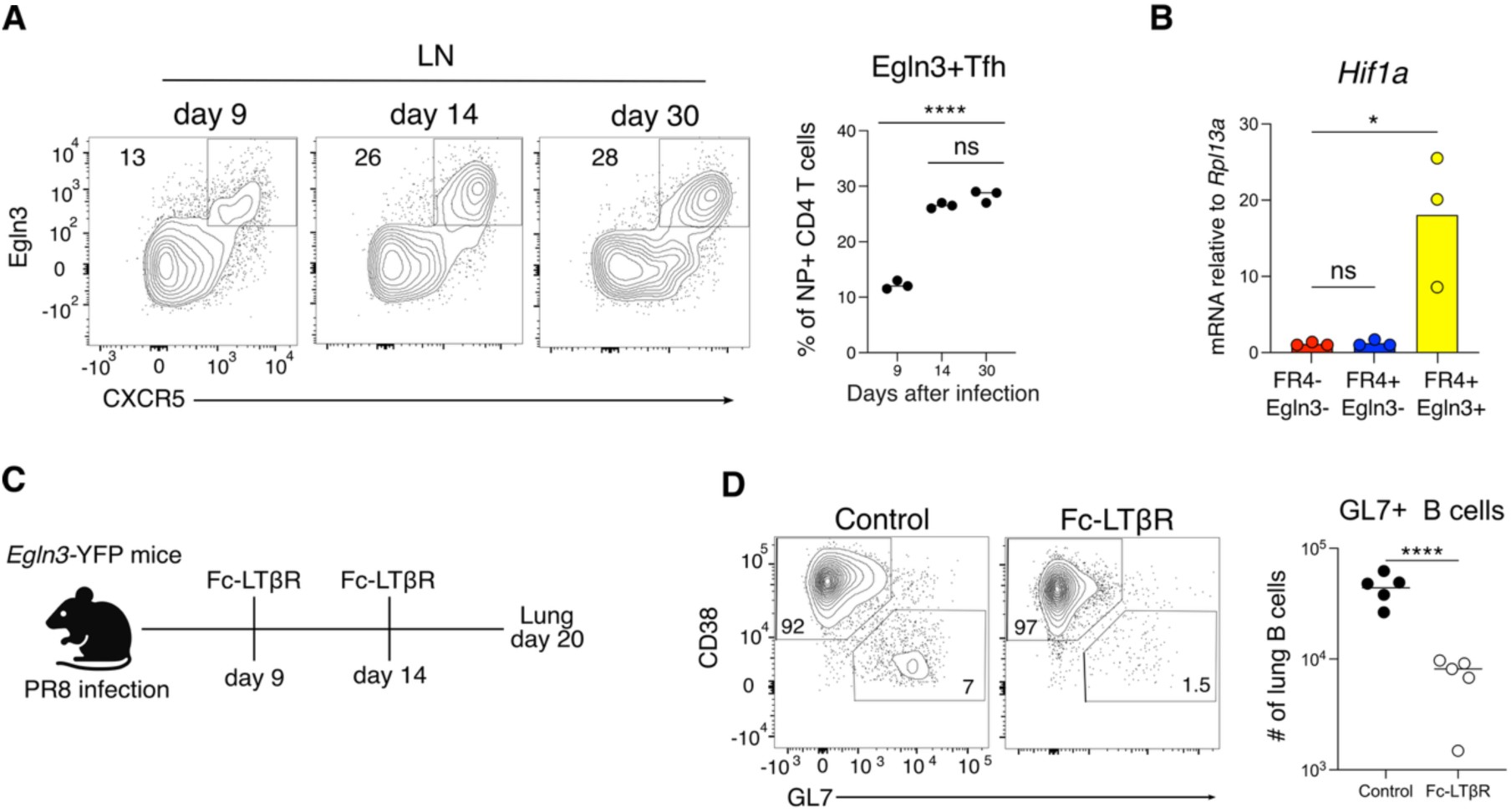
Lung CD4 T cells increase HIF-1α activity during influenza infection. (A) Representative flow cytometry plot and frequency of LN NP+ Tfh cells from Egln3-YFP mice at 9, 14, and 30 days post-PR8 infection (mean ± SD, *n =* 3 mice). (B) Hif1a expression by Taqman qPCR in sorted lung FR4-, FR4+ Egln3-, and FR4+ Egln3+ cells (mean ± SD, *n =* 3 mice). (C) Schematic for TLS disruption by Fc-LTβR during influenza infection. (D) Representative flow cytometry plot and numbers of GL7+ lung B cells after Fc-LTβR administration during influenza infection (mean ± SD, *n =* 5 mice). Statistical significance determined by one-way ANOVA (A,B) and t-test (D). Adjusted p-values: *p<0.05, **p<0.005, ***p<0.0005, ****p<0.0001, ns: not significant. Data representative of two independent experiments.

**Figure S3.**
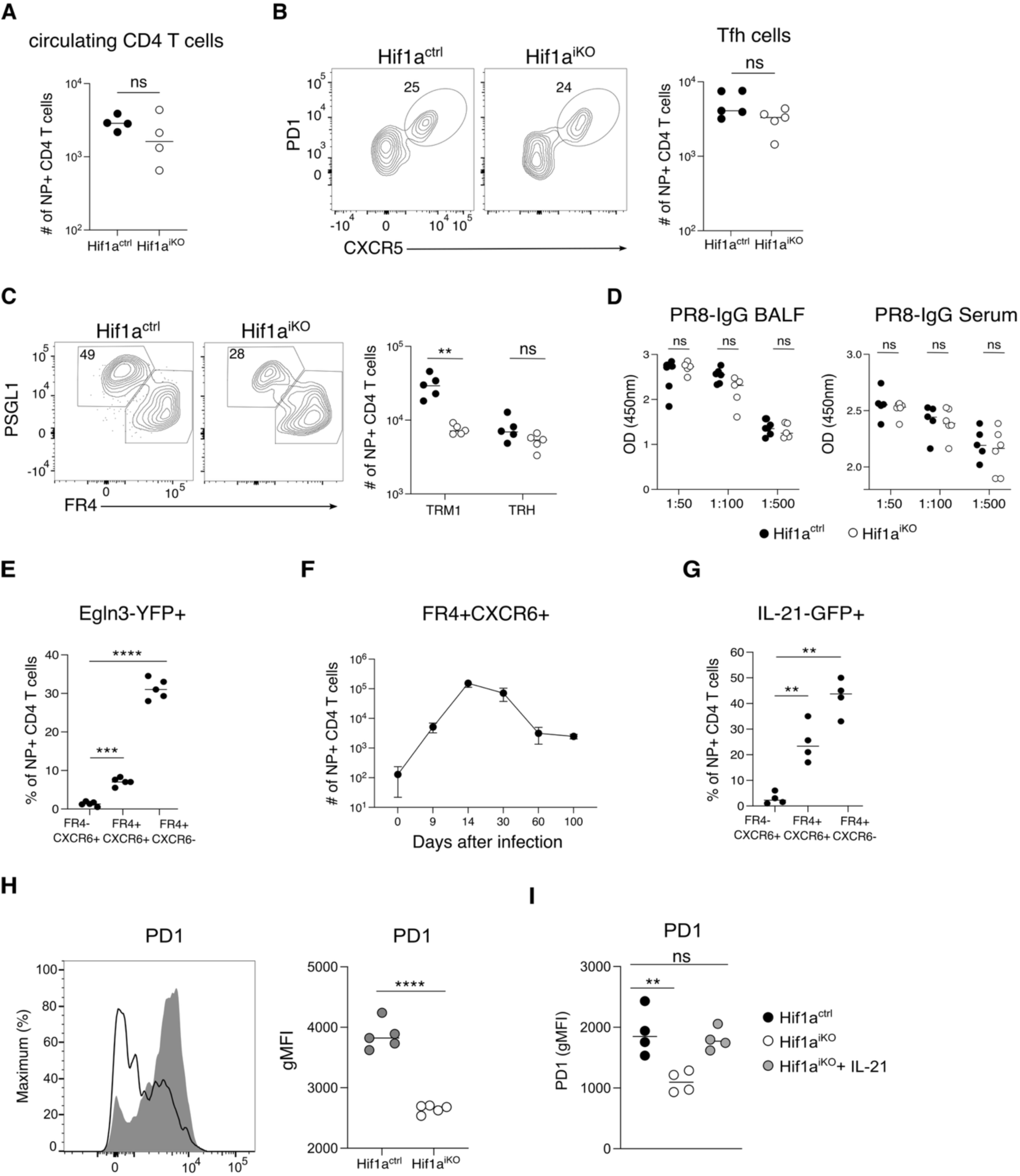
HIF-1α+ CD4 T cells produce IL-21 and support T cell residency in the lung. (A) Numbers of lung circulating NP+ CD4 T cells (mean ± SD, *n =* 4 mice). (B) Representative plots and LN NP+ Tfh cell frequencies in Hif1a^ctrl^ and Hif1a^iKO^ mice (mean ± SD, *n =* 5 mice). (C) Representative plots and NP+ TRM1/TRH cell numbers (mean ± SD, *n =* 5 mice). (D) PR8-specific IgG ELISA from BALF and serum (mean ± SD, *n =* 5-6 mice). (E) Frequency of Egln3-YFP+ NP+ CD4 T cell subsets (mean ± SD, *n =* 5 mice). (F) Time course of FR4+CXCR6+ CD4 T cells post-PR8 infection (mean ± SD, *n =* 4 mice). (G) Frequency of IL-21-GFP+ in NP+ CD4 T cell subsets from IL-21-GFP mice (mean ± SD, *n =* 5-6 mice). (H, I) Histogram and quantification of PD1 expression (gMFI) in NP-specific CD4 T cells using Hif1a^ctrl^ and Hif1a^iKO^ mice and following intratracheal IL-21 treatment (days 14–20) (mean ± SD, *n =* 4-5 mice). Statistical significance determined by t-test (A, B, H), one-way (E, G, I) or two-way ANOVA (C, D). Adjusted p-values: *p<0.05, **p<0.005, ***p<0.0005, ****p<0.0001, ns: not significant. Data representative of two independent experiments.

**Figure S4.**
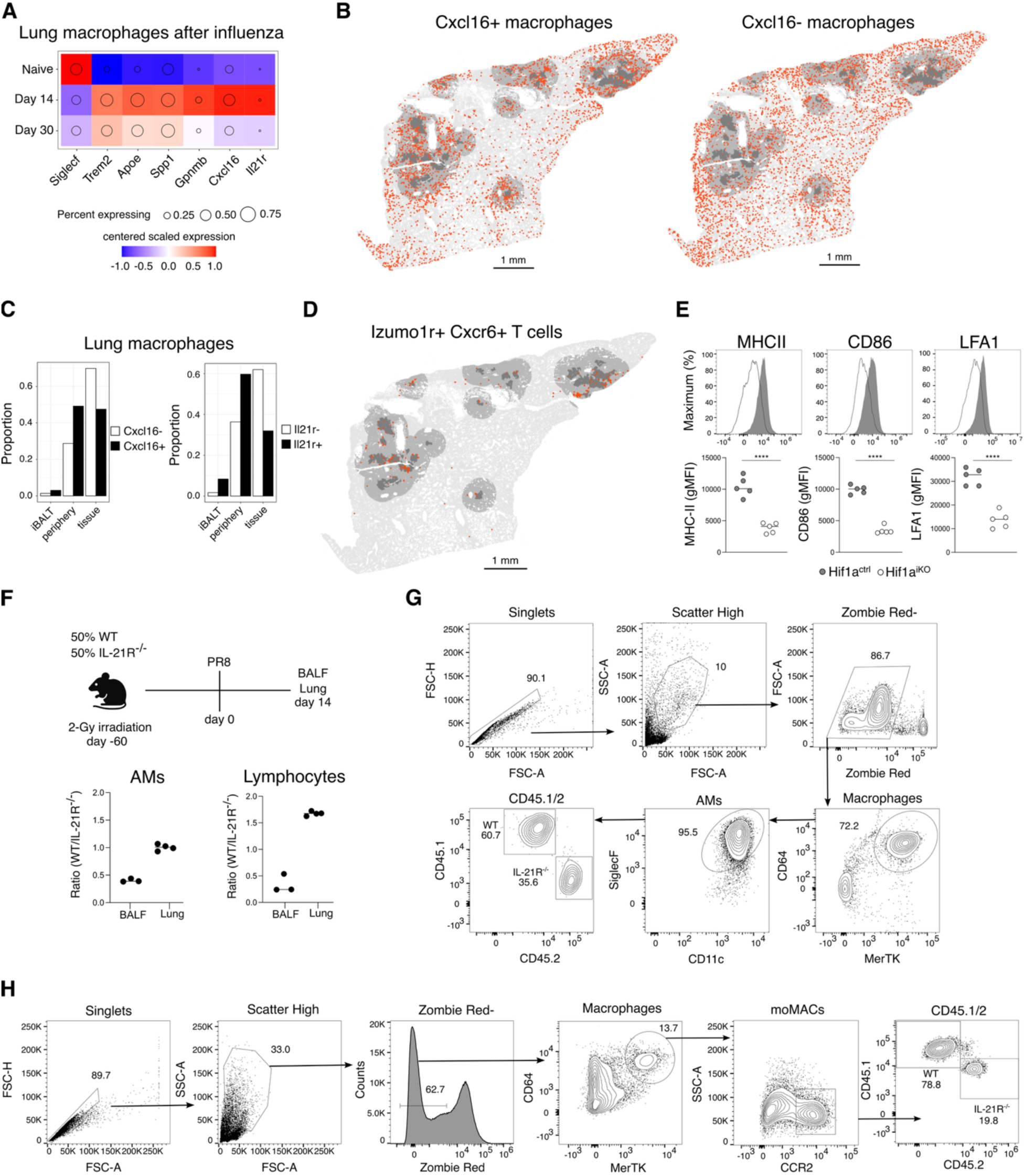
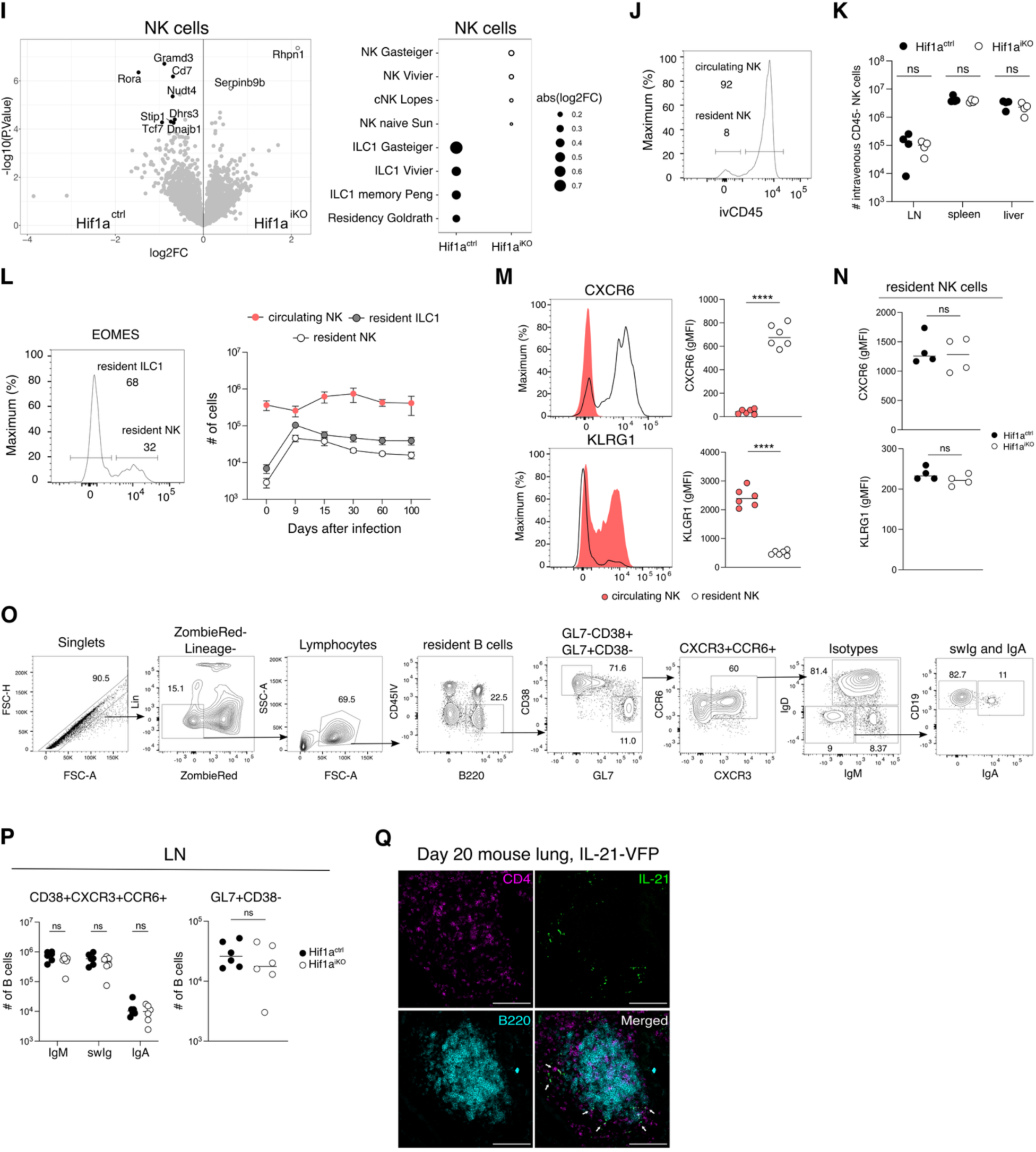
HIF-1α+ CD4 T cells orchestrate the lung immune response during influenza. (A) Centered, scaled, normalized, mean gene expression in lung macrophages over time post influenza. (B–D) Xenium data, day 14 post-influenza: (B) Spatial location of Cxcl16+ vs. Cxcl16-macrophages; (C) Proportion of macrophage phenotypes by region; (D) Spatial location of Izumo1r+Cxcr6+ T cells (*n =* 1 mouse). (E) Flow cytometry and quantification of alveolar macrophages (AMs) from Hif1a^ctrl^ and Hif1a^iKO^ lungs. (F) Chimera schematic (for Figure 4) and CD45.1+ (WT) vs. CD45.2+ (IL-21R⁻/⁻) cell ratios in BALF and lung AMs and lymphocytes at steady state. (G–H) Gating strategies for BALF AMs (G) and lung monocyte-derived macrophages and lymphocytes (H). (I) scRNA-seq from Hif1a^ctrl^ and Hif1a^iKO^ lungs: pseudobulk differential gene expression in NK cells and gene set enrichment at day 30 post-infection (log2FC, *n =* 3 mice). (J) Flow cytometry of IV-CD45+ (circulating) vs. CD45- (resident) NK cells (mean ± SD, *n =* 4 mice). (K) Resident NK counts in LN, spleen, liver (mean ± SD, *n =* 4 mice). (L) Lung ILC1 (EOMES⁻) and conventional NK (EOMES⁺) cells over time (mean ± SD, *n =* 5 mice). (M) CXCR6 and KLRG1 expression (gMFI) in circulating vs. resident NK cells in wild type mice (mean ± SD, *n =* 5 mice). (N) CXCR6 and KLRG1 expression (gMFI) in resident NK cells from Hif1a^ctrl^ and Hif1a^iKO^ mice (mean ± SD, *n =* 5 mice). (O) Gating for lung and LN B cells at day 30 post-infection. (P) Numbers of LN CD38+CXCR3+CCR6+ and GL7+CD38-B cells in Hif1a^ctrl^ and Hif1a^iKO^ (mean ± SD, *n =* 6 mice). (Q) Confocal image of iBALT with IL-21-VFP+ CD4 T cell. Bars, 100µm (mean ± SD, *n =* 3 mice). Statistical significance determined by t-test (B, C, D, M, N, P); one and two-way ANOVA (N, Q). Adjusted p-values: ***p<0.0005; ns: not significant. Data representative of two independent experiments.

**Figure S5.**
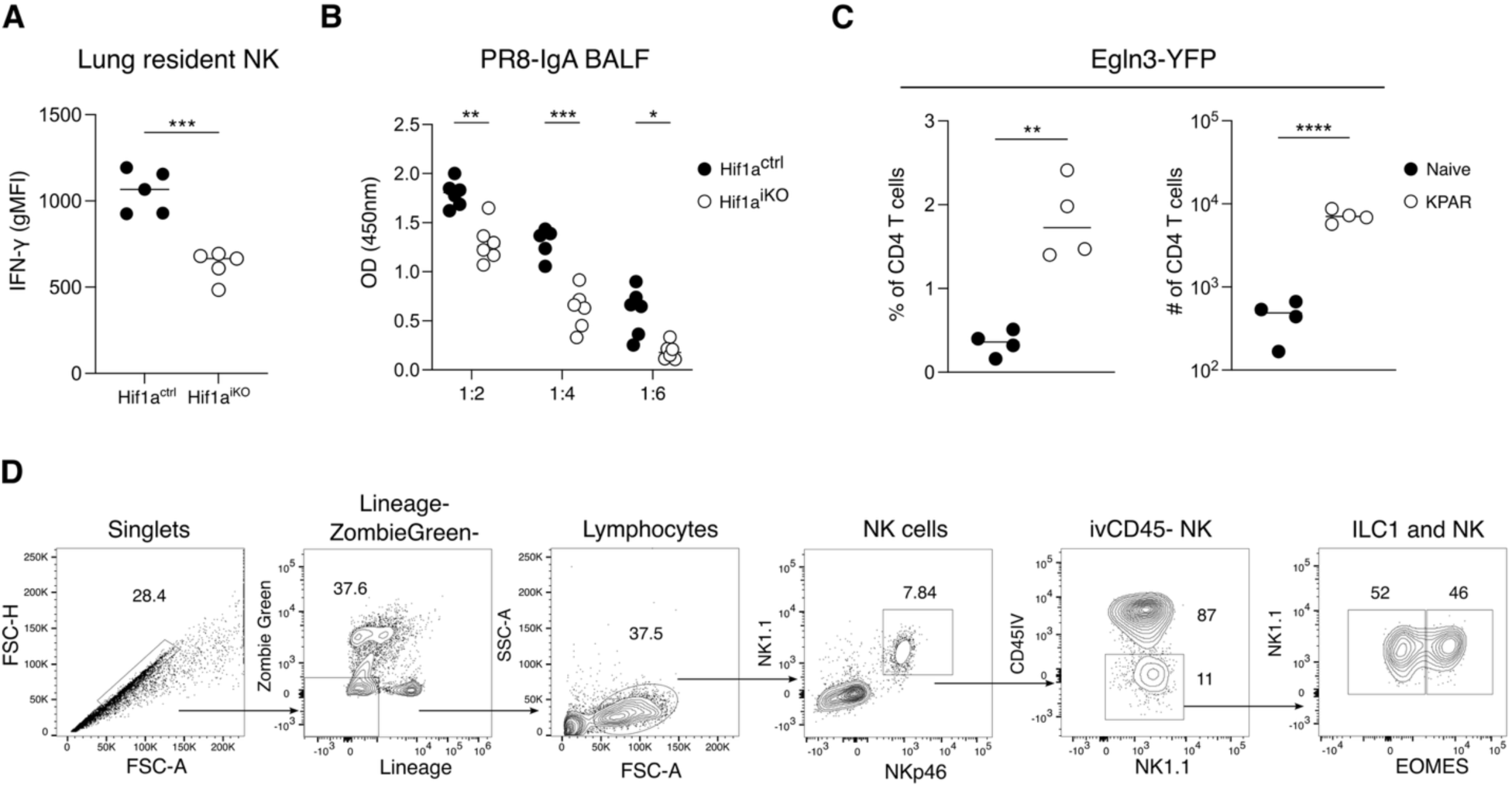
HIF-1α+ CD4 T cells coordinate mucosal immunity to infectious challenge and tumor. (A) IFN-γ expression in resident NK cells (mean ± SD, *n =* 5 mice). (B) PR8 specifc IgA in BALF from Hif1a^ctrl^ and Hif1a^iKO^ mice infected with X31 and followed by PR8 challenge (mean ± SD, *n =* 5 mice). (C) Frequency and numbers of Egln3-YFP+ lung resident CD4 T cells (mean ± SD, *n =* 4 mice). (D) Gating strategy for lung resident ILC1 in KPAR model (mean ± SD, *n =* 5 mice). Statistical analyses were performed using t-tests (A,C) and one-way ANOVA (B). Adjusted p-values: *p<0.05, **p<0.005, ***p<0.0005, ****p<0.0001. Data representative of two independent experiments.

