## Supplemental figures for "HIF-1α+ CD4 T cells coordinate a tissue resident immune cell network in the lung"

### Figure S1

A

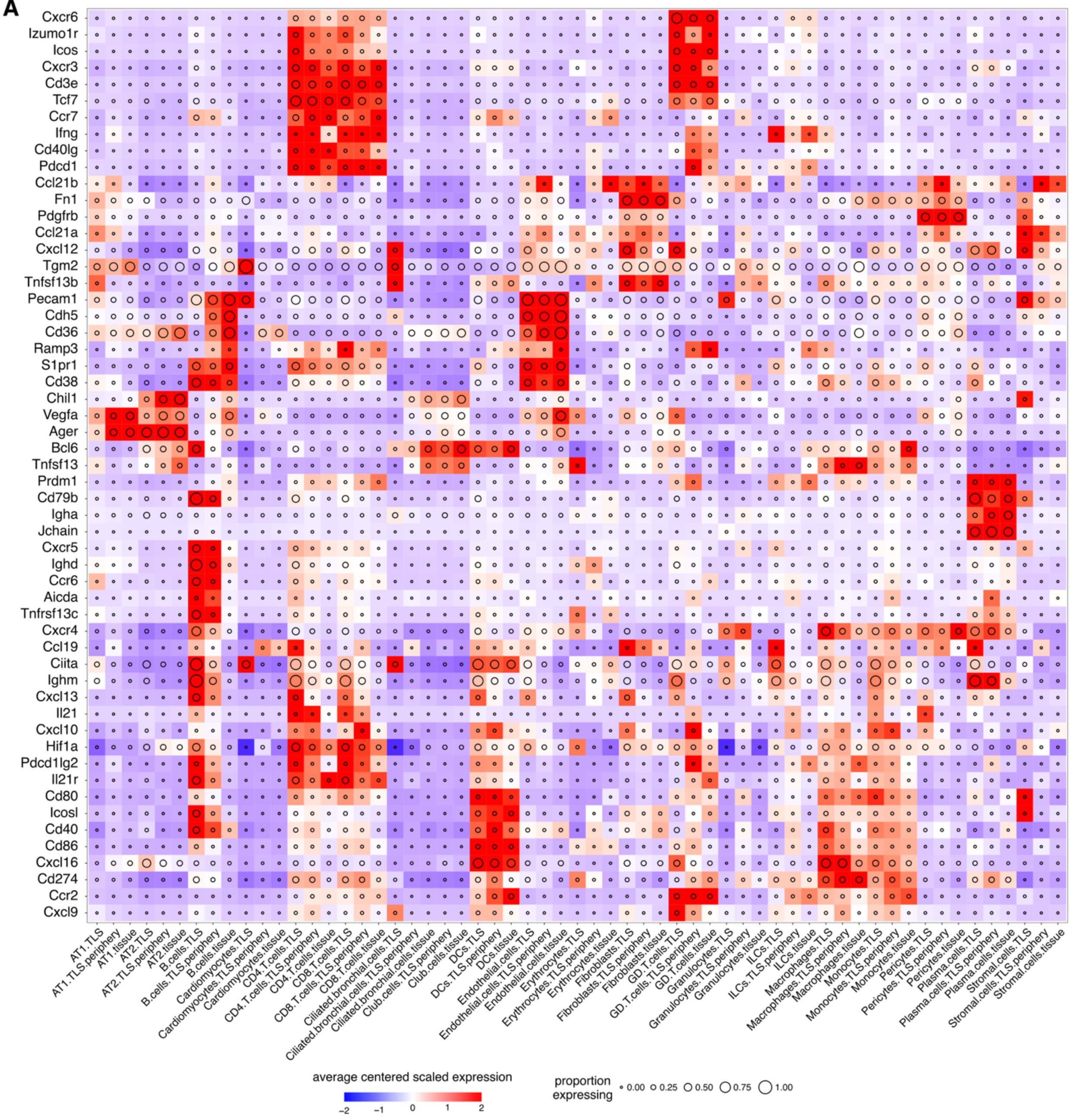
Cxcr6
Izumo1r
Icos
Cxcr3
Cd3e
Tcf7
Ccr7
Ifng
Cd40lg
Pdcd1
Ccl21b
Fn1
Pdgfrb
Ccl21a
Cxcl12
Tgm2
Tnfsf13b
Pecam1
Cdh5
Cd36
Ramp3
S1pr1
Cd38
Chil1
Vegfa
Ager
Bcl6
Tnfsf13
Prdm1
Cd79b
Igha
Jchain
Cxcr5
Ighd
Ccr6
Aicda
Tnfrsf13c
Cxcr4
Ccl19
Ciita
Ighm
Cxcl13
Il21
Cxcl10
Hif1a
Pdcd1lg2
Il21r
Cd80
Icosl
Cd40
Cd86
Cxcl16
Cd274
Ccr2
Cxcl9
AT1.TLS
AT1.TLS.periphery
AT1.tissue
AT2.TLS
AT2.TLS.periphery
AT2.tissue
B.cells.TLS
B.cells.TLS.periphery
B.cells.tissue
Cardiomyocytes.TLS
Cardiomyocytes.TLS.periphery
Cardiomyocytes.tissue
CD4.T.cells.TLS
CD4.T.cells.TLS.periphery
CD4.T.cells.tissue
CD8.T.cells.TLS
CD8.T.cells.TLS.periphery
CD8.T.cells.tissue
Ciliated.bronchial.cells.TLS
Ciliated.bronchial.cells.TLS.periphery
Ciliated.bronchial.cells.tissue
Club.cells.TLS.periphery
Club.cells.tissue
DCs.TLS
DCs.TLS.periphery
DCs.tissue
Endothelial.cells.TLS
Endothelial.cells.TLS.periphery
Endothelial.cells.tissue
Erythrocytes.TLS
Erythrocytes.TLS.periphery
Erythrocytes.tissue
Fibroblasts.TLS
Fibroblasts.TLS.periphery
Fibroblasts.tissue
GD.T.cells.TLS
GD.T.cells.TLS.periphery
GD.T.cells.tissue
Granulocytes.TLS
Granulocytes.TLS.periphery
Granulocytes.tissue
ILCs.TLS
ILCs.TLS.periphery
ILCs.tissue
Macrophages.TLS
Macrophages.TLS.periphery
Macrophages.tissue
Monocytes.TLS
Monocytes.TLS.periphery
Monocytes.tissue
Pericytes.TLS
Pericytes.TLS.periphery
Pericytes.tissue
Plasma.cells.TLS
Plasma.cells.TLS.periphery
Plasma.cells.tissue
Stromal.cells.TLS
Stromal.cells.TLS.periphery
Stromal.cells.tissue
average centered scaled expression
−2
−1
0
1
2
proportion expressing
0.00
0.25
0.50
0.75
1.00

**Figure S1**

**B**

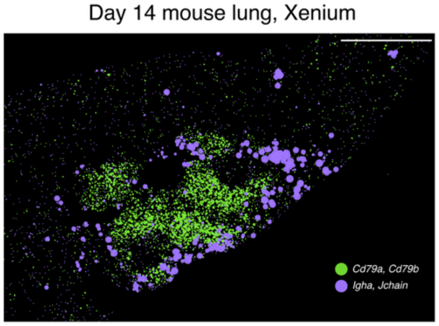


**C**

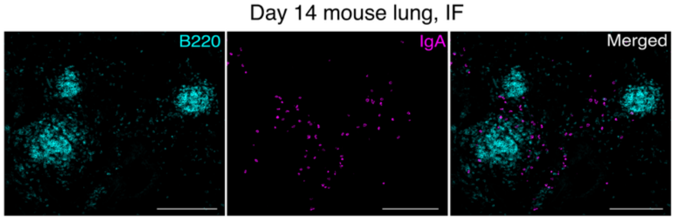


**Figure S1. Spatially structured immune cell interactions in lung TLS.**

(A) Centered, scaled normalized expression of Xenium data of mouse lung day 14 post-influenza. Expression averaged per cell type and tissue location (*n* = 1 mouse).

(B) Representative iBALT region from day 14 lung as displayed in Xenium Explorer. Bars, 500µm.

(C) Immunofluorescence (IF) showing inducible iBALTs (cyan) containing IgA$^+$ cells (magenta) 14 days after influenza infection. Bars, 200µm (*n* = 3 mice).

**Figure S2**

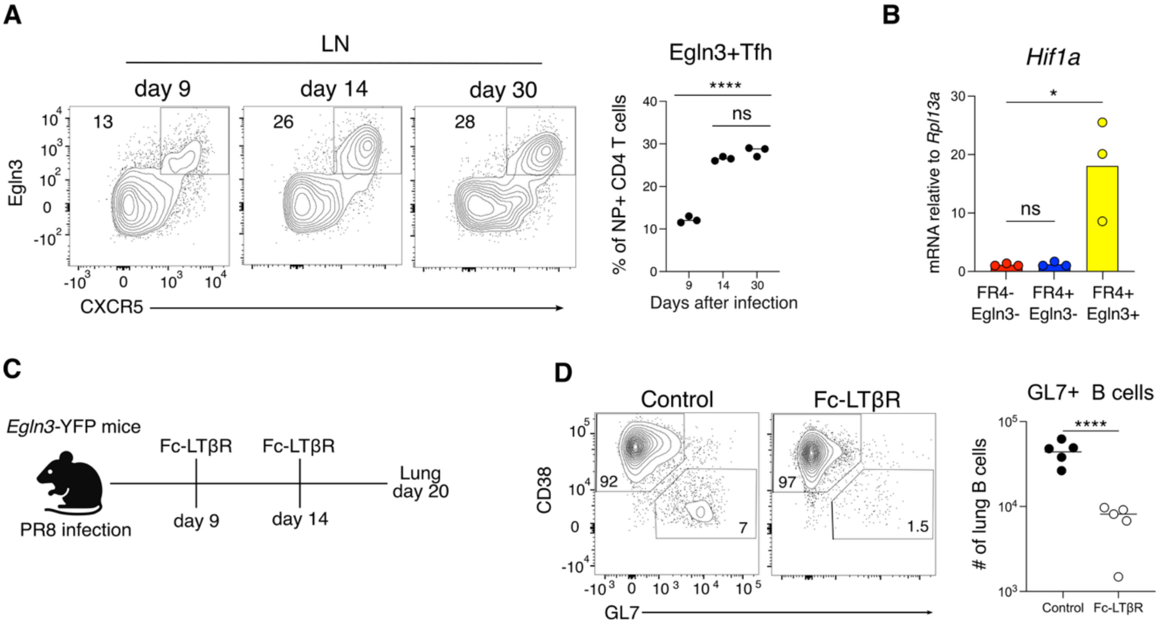


**Figure S2. Lung CD4 T cells increase HIF-1α activity during influenza infection.**

(A) Representative flow cytometry plot and frequency of LN NP+ Tfh cells from Egln3-YFP mice at 9, 14, and 30 days post-PR8 infection (mean ± SD, *n* = 3 mice).

(B) Hif1a expression by Taqman qPCR in sorted lung FR4-, FR4+ Egln3-, and FR4+ Egln3+ cells (mean ± SD, *n* = 3 mice).

(C) Schematic for TLS disruption by Fc-LTβR during influenza infection.

(D) Representative flow cytometry plot and numbers of GL7+ lung B cells after Fc-LTβR administration during influenza infection (mean ± SD, *n* = 5 mice).

Statistical significance determined by one-way ANOVA (A, B) and t-test (D). Adjusted p-values: *p<0.05, **p<0.005, ***p<0.0005, ****p<0.0001, ns: not significant. Data representative of two independent experiments.

**Figure S3**

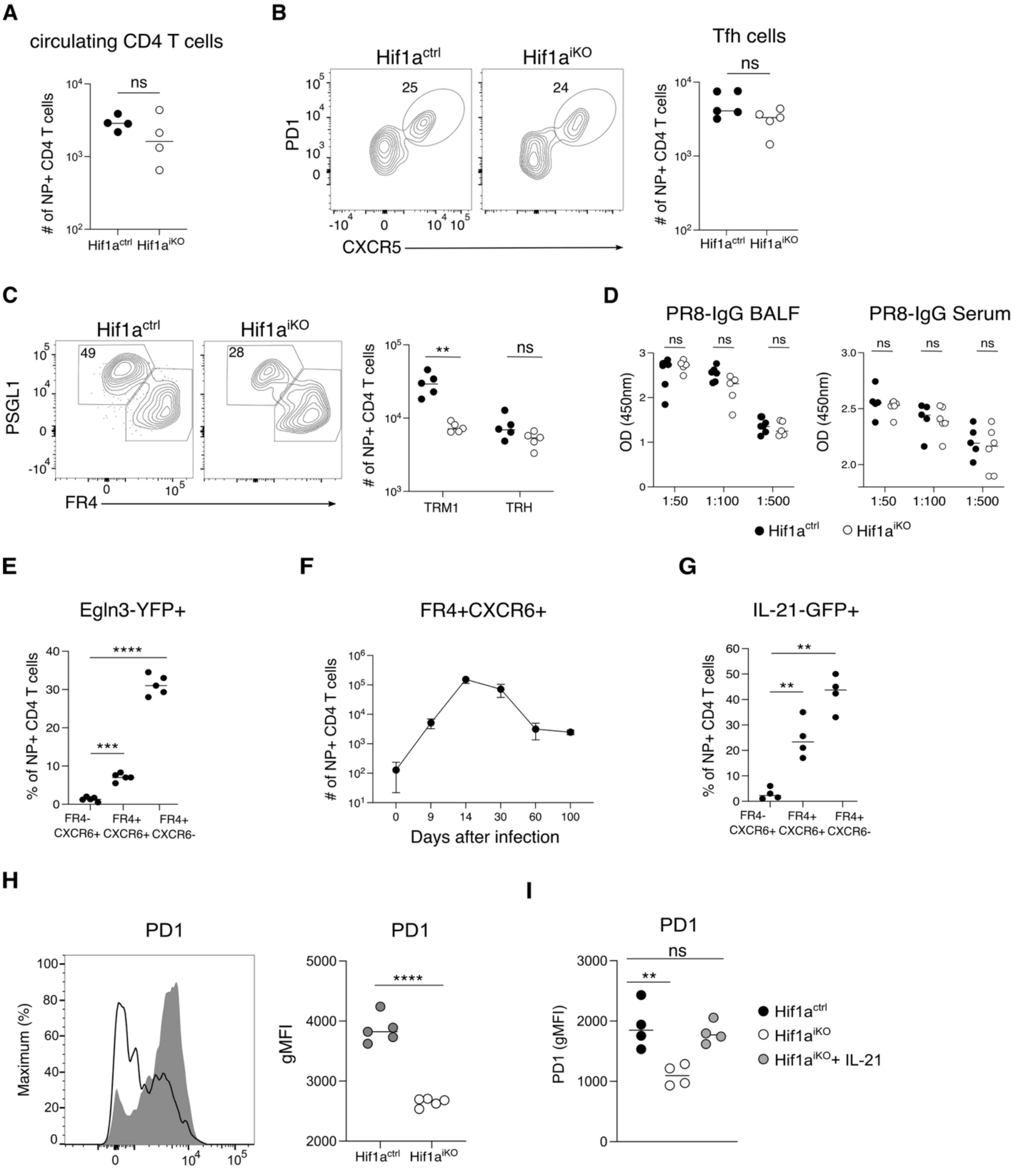


**Figure S3. HIF-1α+ CD4 T cells produce IL-21 and support T cell residency in the lung.**

(A) Numbers of lung circulating NP+ CD4 T cells (mean ± SD, *n* = 4 mice).

(B) Representative plots and LN NP+ Tfh cell frequencies in $Hif1a^{ctrl}$ and $Hif1a^{iKO}$ mice (mean ± SD, *n* = 5 mice).

(C) Representative plots and NP+ TRM1/TRH cell numbers (mean ± SD, *n* = 5 mice).

(D) PR8-specific IgG ELISA from BALF and serum (mean ± SD, *n* = 5-6 mice).

(E) Frequency of Egln3-YFP+ NP+ CD4 T cell subsets (mean ± SD, *n* = 5 mice).

(F) Time course of FR4+CXCR6+ CD4 T cells post-PR8 infection (mean ± SD, *n* = 4 mice).

(G) Frequency of IL-21-GFP+ in NP+ CD4 T cell subsets from IL-21-GFP mice (mean ± SD, *n* = 5-6 mice).

(H, I) Histogram and quantification of PD1 expression (gMFI) in NP-specific CD4 T cells using $Hif1a^{ctrl}$ and $Hif1a^{iKO}$ mice and following intratracheal IL-21 treatment (days 14–20) (mean ± SD, *n* = 4-5 mice).

Statistical significance determined by t-test (A, B, H), one-way (E, G, I) or two-way ANOVA (C, D). Adjusted p-values: *p<0.05, **p<0.005, ***p<0.0005, ****p<0.0001, ns: not significant. Data representative of two independent experiments.

**Figure S4**

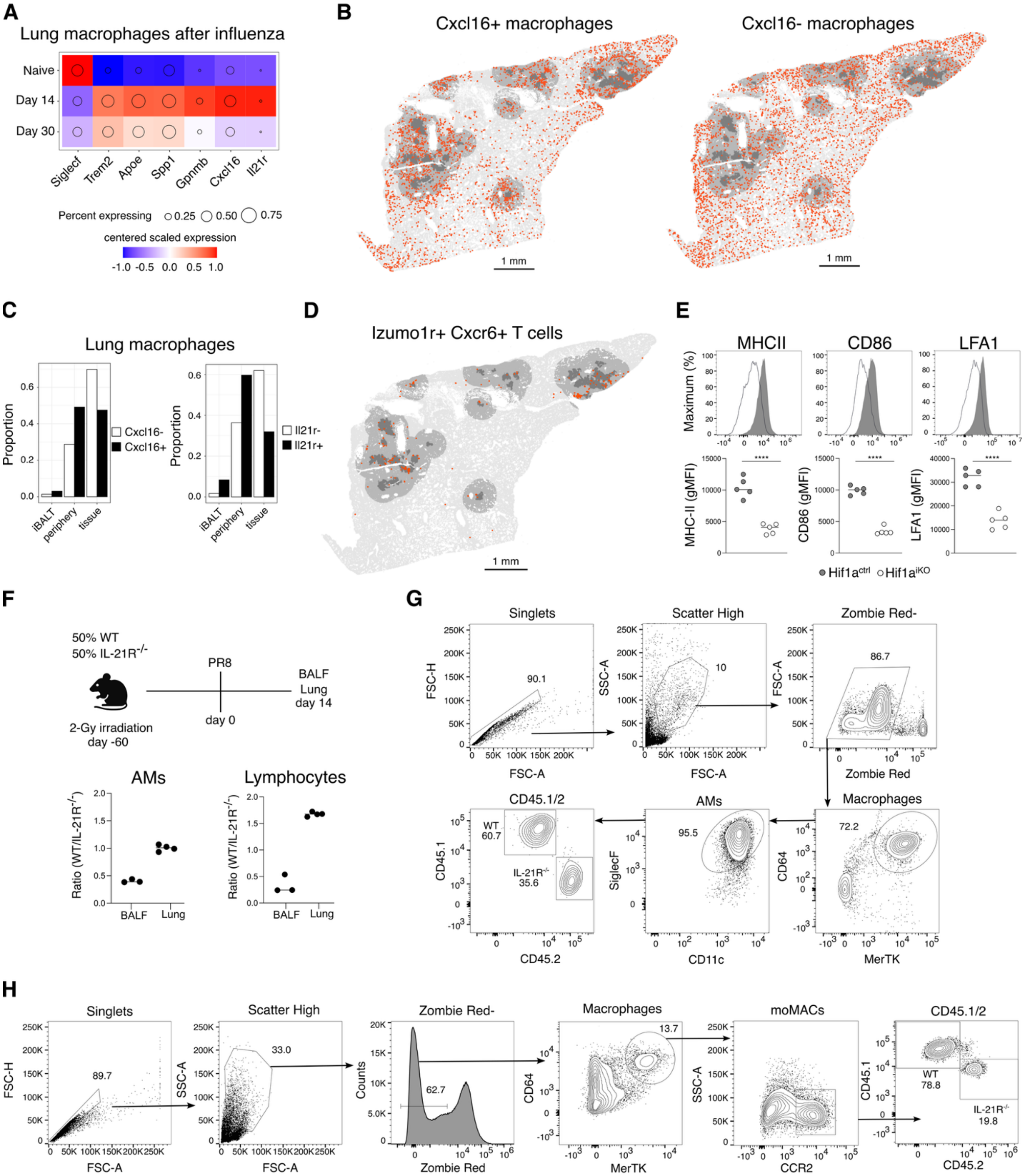
A
Lung macrophages after influenza
Naive
Day 14
Day 30
Siglecf
Trem2
Apoe
Spp1
Gpnmb
Cxcl16
Il21r
Percent expressing
0.25
0.50
0.75
centered scaled expression
-1.0 -0.5 0.0 0.5 1.0
B
Cxcl16+ macrophages
Cxcl16- macrophages
1 mm
C
Lung macrophages
Proportion
Cxcl16-
Cxcl16+
Il21r-
Il21r+
iBALT
periphery
tissue
D
Izumo1r+ Cxcr6+ T cells
1 mm
E
MHCII
CD86
LFA1
Maximum (%)
MHC-II (gMFI)
CD86 (gMFI)
LFA1 (gMFI)
****
Hif1a ctrl
Hif1a iKO
F
50% WT
50% IL-21R-/-
PR8
BALF
Lung
day 14
2-Gy irradiation
day -60
day 0
AMs
Lymphocytes
Ratio (WT/IL-21R-/-)
BALF
Lung
G
Singlets
Scatter High
Zombie Red-
FSC-H
FSC-A
SSC-A
90.1
10
86.7
Zombie Red
CD45.1/2
AMs
Macrophages
WT
60.7
IL-21R-/-
35.6
95.5
72.2
CD45.1
CD45.2
SiglecF
CD11c
CD64
MerTK
H
Singlets
Scatter High
Zombie Red-
Macrophages
moMACs
CD45.1/2
89.7
33.0
62.7
13.7
Counts
Zombie Red
CD64
MerTK
SSC-A
CCR2
CD45.1
CD45.2
WT
78.8
IL-21R-/-
19.8

**Figure S4**

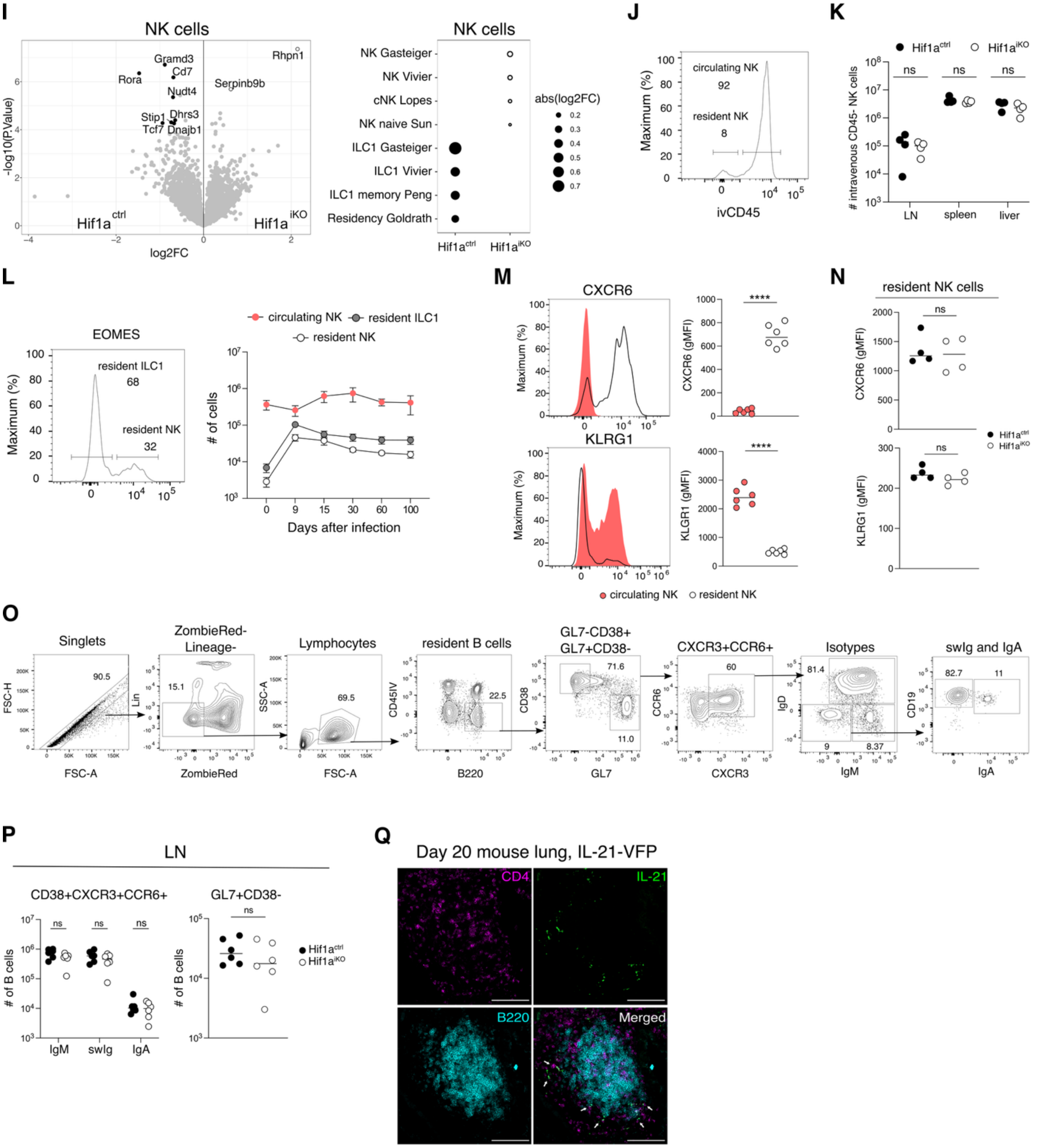


**Figure S4. HIF-1α+ CD4 T cells orchestrate the lung immune response during influenza.**

(A) Centered, scaled, normalized, mean gene expression in lung macrophages over time post influenza.

(B–D) Xenium data, day 14 post-influenza: (B) Spatial location of Cxcl16+ vs. Cxcl16- macrophages; (C) Proportion of macrophage phenotypes by region; (D) Spatial location of Izumo1r+Cxcr6+ T cells (*n* = 1 mouse).

(E) Flow cytometry and quantification of alveolar macrophages (AMs) from $Hif1a^{ctrl}$ and $Hif1a^{iKO}$ lungs.

(F) Chimera schematic (for Figure 4) and CD45.1+ (WT) vs. CD45.2+ ($IL\text{-}21R^{-}/^{-}$) cell ratios in BALF and lung AMs and lymphocytes at steady state.

(G–H) Gating strategies for BALF AMs (G) and lung monocyte-derived macrophages and lymphocytes (H).

(I) scRNA-seq from $Hif1a^{ctrl}$ and $Hif1a^{iKO}$ lungs: pseudobulk differential gene expression in NK cells and gene set enrichment at day 30 post-infection (log2FC, *n* = 3 mice).

(J) Flow cytometry of IV-CD45+ (circulating) vs. CD45- (resident) NK cells (mean ± SD, *n* = 4 mice).

(K) Resident NK counts in LN, spleen, liver (mean ± SD, *n* = 4 mice).

(L) Lung ILC1 ($EOMES^{-}$) and conventional NK ($EOMES^{+}$) cells over time (mean ± SD, *n* = 5 mice).

(M) CXCR6 and KLRG1 expression (gMFI) in circulating vs. resident NK cells in wild type mice (mean ± SD, *n* = 5 mice).

(N) CXCR6 and KLRG1 expression (gMFI) in resident NK cells from $Hif1a^{ctrl}$ and $Hif1a^{iKO}$ mice (mean ± SD, *n* = 5 mice).

(O) Gating for lung and LN B cells at day 30 post-infection.

(P) Numbers of LN CD38+CXCR3+CCR6+ and GL7+CD38- B cells in $Hif1a^{ctrl}$ and $Hif1a^{iKO}$ (mean ± SD, *n* = 6 mice).

(Q) Confocal image of iBALT with IL-21-VFP+ CD4 T cell. Bars, 100μm (mean ± SD, *n* = 3 mice).

Statistical significance determined by t-test (B, C, D, M, N, P); one and two-way ANOVA (N, Q). Adjusted p-values: ***p<0.0005; ns: not significant. Data representative of two independent experiments.

#### Figure S5

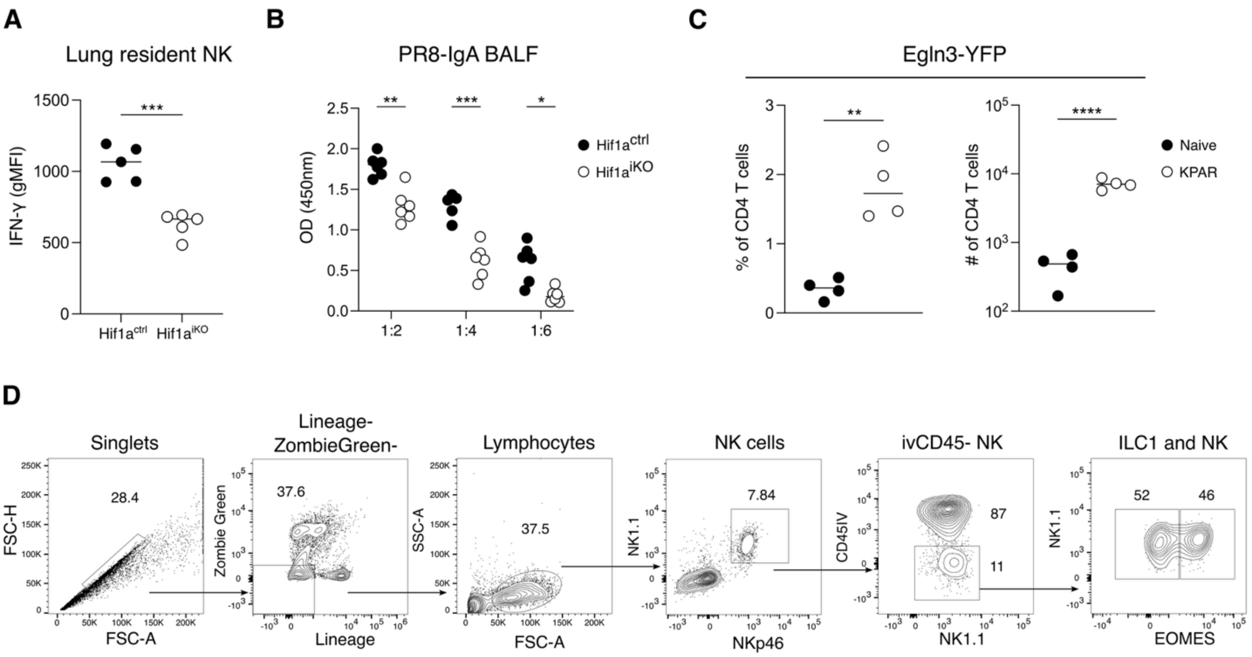


**Figure S5. HIF-1α+ CD4 T cells coordinate mucosal immunity to infectious challenge and tumor.**

(A) IFN-γ expression in resident NK cells (mean ± SD, *n* = 5 mice).

(B) PR8 specifc IgA in BALF from Hif1a$^{ctrl}$ and Hif1a$^{iKO}$ mice infected with X31 and followed by PR8 challenge (mean ± SD, *n* = 5 mice).

(C) Frequency and numbers of Egln3-YFP+ lung resident CD4 T cells (mean ± SD, *n* = 4 mice).

(D) Gating strategy for lung resident ILC1 in KPAR model (mean ± SD, *n* = 5 mice).

Statistical analyses were performed using t-tests (A, C) and one-way ANOVA (B). Adjusted p-values: *p<0.05, **p<0.005, ***p<0.0005, ****p<0.0001. Data representative of two independent experiments.
